# Temporal regulation of a spatial patterning factor in *Drosophila* neurogenesis

**DOI:** 10.64898/2026.08.11.744200

**Authors:** Rose Coyne, Fahad Kamulegeya, Cathleen Lake, Raghuvanshi Rajesh, McKenzie Treese, Yen-Chung Chen, Benjamin Troutwine, Julia Zeitlinger, Mehmet Neset Özel

**Affiliations:** Stowers Institute for Medical Research, Kansas City, MO 64110, USA; Department of Biology, New York University, New York, NY 10003, USA

## Abstract

A central question in neurobiology is how the transient programs that pattern neural progenitors are translated into the enormous, stable diversity of neuronal types. Spatial and temporal cues act only briefly, yet each neuron’s identity is defined and maintained for life by terminal selector transcription factors (TFs). How a neuron’s developmental origin is read out into a particular selector code remains poorly understood. Some current models propose that spatial and temporal origins are inherited independently through separate selectors. We show instead that, in the *Drosophila* optic lobe, the same selector can be activated by different patterning axes through physically distinct enhancers, even within the same lineage. Visual system homeobox (Vsx1) spatially patterns a central neuroepithelial domain and later acts as a terminal selector in dozens of neuronal types, most originating exclusively from that domain. However, in Dm2 neurons that are produced from every domain, it is regulated not by neuroepithelial Vsx1 but by the neuroblast temporal TF BarH1, through an enhancer distinct from its domain-specific ones. Combining *in vivo* reporters with sequence-to-accessibility deep-learning models, we identify and disrupt the key binding sites in this enhancer, impairing its Dm2-specific activity. Reciprocally, the temporal TF Homeobrain (Hbn) acts as a terminal selector in the related neuron Mi21 independently of its neuroblast temporal window: its expression in these late-born neurons is instead placed under dorsoventral spatial control. Patterning inputs therefore need not be partitioned across separate selectors but converge combinatorially on the modular enhancers of shared ones, revealing a cis-regulatory logic that re-encodes this limited set of inputs into vast neuronal diversity.

## Introduction

The remarkable functional capacity of a nervous system rests on the diversity of the neurons that compose it. The scale of this diversity poses a fundamental problem in developmental genetics: how can a small, finite pool of neural stem cells reproducibly give rise to thousands of distinct cell types^1,2^, each with its own morphology, connectivity, and physiology? It is useful to frame the question in terms of the two genetic programs that must ultimately be reconciled: the patterning programs that diversify progenitors and assign each neuron a developmental origin, and the identity programs that establish and maintain the specific properties of each differentiated neuron. How the former is translated into the latter, i.e. how exactly a neuron’s developmental origin is read out as its identity, remains a central and largely unresolved question in neurobiology.

The *Drosophila* optic lobe is an ideal model system in which to dissect this translation. Its repetitive, retinotopic architecture^3^ packs hundreds of well-defined cell types into a structure whose complete EM connectome has now been traced^4,5^, while single-cell atlases resolve on the order of 250 molecular cell types that map cleanly onto these morphological classes^6,7^. Most optic lobe-intrinsic neurons are generated during late larval to early pupal stages by the outer proliferation center (OPC)^8^. In the medial OPC, which mostly generates neurons of the medulla neuropil, neuroepithelial (NE) cells develop into neuroblasts (NB) that divide asymmetrically to self-renew and bud off ganglion mother cells (GMC); each GMC then divides once more to produce two postmitotic cells^9,10^. This process occurs gradually in a mediolateral wave in the developing optic lobes^11^, such that different copies of the same cell type are produced continuously over a 2–3-day period. This is a unique feature of the *Drosophila* visual system, and it previously allowed us to reconstruct complete trajectories, from NE to young neurons, within a single snapshot taken at late L3^12^ or P0^7^ stage using single-cell genomics. As a result, the developmental origin of nearly every OPC neuron has been mapped^13^, providing well-defined coordinates against which to test how identity is specified.

Neuronal diversity in the OPC is instructed by the intersection of at least three patterning axes. Spatially, the NE is subdivided into discrete domains by the transcription factors (TFs) Vsx1, Optix, Rx, Disco and Spalt, as well as by the signaling proteins Wg (Wnt) and Dpp (BMP)^14-^16. Then, each NB advances through a sequence of at least twelve temporal TFs, so that neurons born at successive times inherit different patterning inputs^12,17,18^. Finally, the Notch-dependent split between sister neurons from the GMC division provides a third, binary axis^18^. The combinatorial intersection of spatial domain, temporal window, and Notch status can account for the origin of essentially every medulla cell type^13^. Recently, a fourth axis has been proposed whereby the NE is also subjected to temporal patterning by RNA-binding proteins Imp and Syp^19^, further diversifying the patterning inputs. A recurring and consequential observation, however, is that the spatial and temporal factors that pattern progenitors are, with very few exceptions, not maintained in the neurons they help to specify.

This last point lies at the heart of how developmental origin becomes cell type identity. Even though neuroblast temporal TFs can often still be detected in their immediate neural progeny^12,20,21^, patterning factors are generally transient and act instead to install a downstream regulatory program in postmitotic neurons^7,22^. These downstream regulators are the terminal selectors (tsTFs): factors continuously expressed in a neuron from birth through adulthood that regulate the effector genes defining its type^23,24^. We previously inferred the putative tsTF codes of nearly all optic lobe neurons and showed that combinations of roughly 10 tsTFs suffice to define each of ∼200 cell types, with targeted changes to these codes driving predictable switches of neuronal identity^25^. Notably, some spatial and temporal patterning TFs also act as terminal selectors in postmitotic neurons. However, neurons that use these TFs do not necessarily descend from the progenitors that were using that same TF as a patterning factor^7,13^, suggesting independent regulation. The key unresolved problem is therefore a regulatory one: *how* the patterning state of each progenitor is translated into faithful, cell-type-specific activation of the correct tsTF combinations through defined enhancers.

Two differing views have emerged on how patterning is organized to control selector expression. Recent studies of the *Drosophila* central brain^26^ and VNC^27^ advance a model in which hemilineage identity (shared spatial origin and Notch status) and temporal (birth-order) patterning operate as independent, orthogonal axes that often govern separate sets of tsTFs in the neuronal progeny. In the optic lobe, by contrast, correlations between developmental origin and selector expression across a large single-cell dataset suggested that most tsTFs are associated not with a single patterning axis but with variable combinations of spatial, temporal, and Notch inputs^13^. This would imply that patterning inputs are often not partitioned across separate selectors. But whether the same selector can genuinely be reached by distinct patterning inputs within the same lineage, and if so through what cis-regulatory logic, has not been demonstrated.

Our recent single-cell multiome (simultaneous RNA + ATAC-seq) atlas of the developing optic lobe offers an inroad into this problem. It revealed that tsTF codes are installed within a brief window in newborn neurons, frequently through enhancers that are not accessible in their progenitors. Notably, when a temporal TF is re-deployed as a tsTF in neurons, it is associated with accessible enhancers entirely distinct from those that are open in neuroblasts^7^. This implies that these genes are independently re-activated in neurons rather than maintained from progenitors, but this has not been demonstrated directly. It is also unclear whether this independence extends beyond temporal TFs. Visual system homeobox 1 (Vsx1) presents the strongest putative case for maintenance: it acts as a spatial patterning factor in a central domain of the OPC NE before being downregulated in NBs^14^, and also as a terminal selector in dozens of optic lobe neuronal types^6,25^. In neurons, *Vsx1* is always co-expressed with *Vsx2,* and the two paralogous genes are located head-to-head within ∼35kb of each other (**Fig. 3a**), likely sharing enhancers. The tight correlation between the Vsx-domain origin and neuronal *Vsx1/2* expression has long made spatial patterning the obvious hypothesis for regulation of these genes.

Here, focusing on three closely-related neurons, we show that Vsx1/2 in Dm2 and Homeobrain (Hbn) in Mi21 act as terminal selectors, each necessary and sufficient to distinguish these cell types from Mi15 neurons. We find that the NE Vsx1 is indeed required for *Vsx1/2* expression in neurons that originate exclusively from the Vsx-domain, but not in Dm2 neurons that are produced from the entire main OPC^15^. We show that *Vsx1/2* expression in Dm2 is instead specified by the neuroblast temporal TF BarH1. It is therefore independent of Vsx1’s role in spatial patterning, even in Dm2 neurons born within the Vsx domain. We find that the spatial and temporal inputs to *Vsx1/2* are routed through physically distinct enhancers, so that the same tsTF can be switched on by different specification factors in different neurons, including those arising from the same stem cell. The converse arrangement also occurs: *hbn*, itself a neuroblast temporal TF, is placed under spatial control when it is expressed as a tsTF in Mi21. Together, these results establish that a single terminal selector gene can be activated independently by different patterning mechanisms, through dedicated cis-regulatory elements. This logic is incompatible with a strict orthogonal partitioning of patterning inputs into separate sets of tsTFs; it argues instead for a modular organization in which distinct enhancers integrate whichever patterning signals are available to assemble each neuron’s unique identity.

## Results

### Vsx1/2 and Hbn instruct the differential identity of Dm2, Mi15 and Mi21 neurons

We first set out to understand how tsTFs in closely related neurons establish cell type identity by investigating a developmentally related group of neurons known as Dm2, Mi15 and Mi21 (**Fig. 1a**), which we collectively refer to as metacluster 24 (M24)^7^. All three of these cell types are Notch^OFF^ (N^OFF^) neurons born during the late temporal windows of OPC neuroblasts^12^. The Dm2 neurons are abundant (one per visual column) and are produced from the entire main OPC while the morphologically similar (but less abundant) Mi15 and Mi21 neurons are made exclusively from dorsal and ventral Optix domains, respectively^13^. They express several tsTFs in common including *Distal-less* (*Dll*), but (among these neurons) Dm2 specifically express *Vsx1/2*, and Mi21 specifically express *hbn* (**Fig. 1b-c**). To be able to visualize and perturb this group of neurons, we generated an *M24-Gal4* line utilizing an ATAC peak^7^ specifically accessible in these three cell types throughout development (**Suppl. Fig. 1**). Its activity in M24 neurons was validated *in vivo*, as it faithfully and exclusively labeled all M24 cell types (with some off-target lamina glia) from P24 to Adult stage, but only Dm2 and Mi15 during neurogenesis at late L3 stage (**Suppl. Fig. 2**).

**Figure 1:**
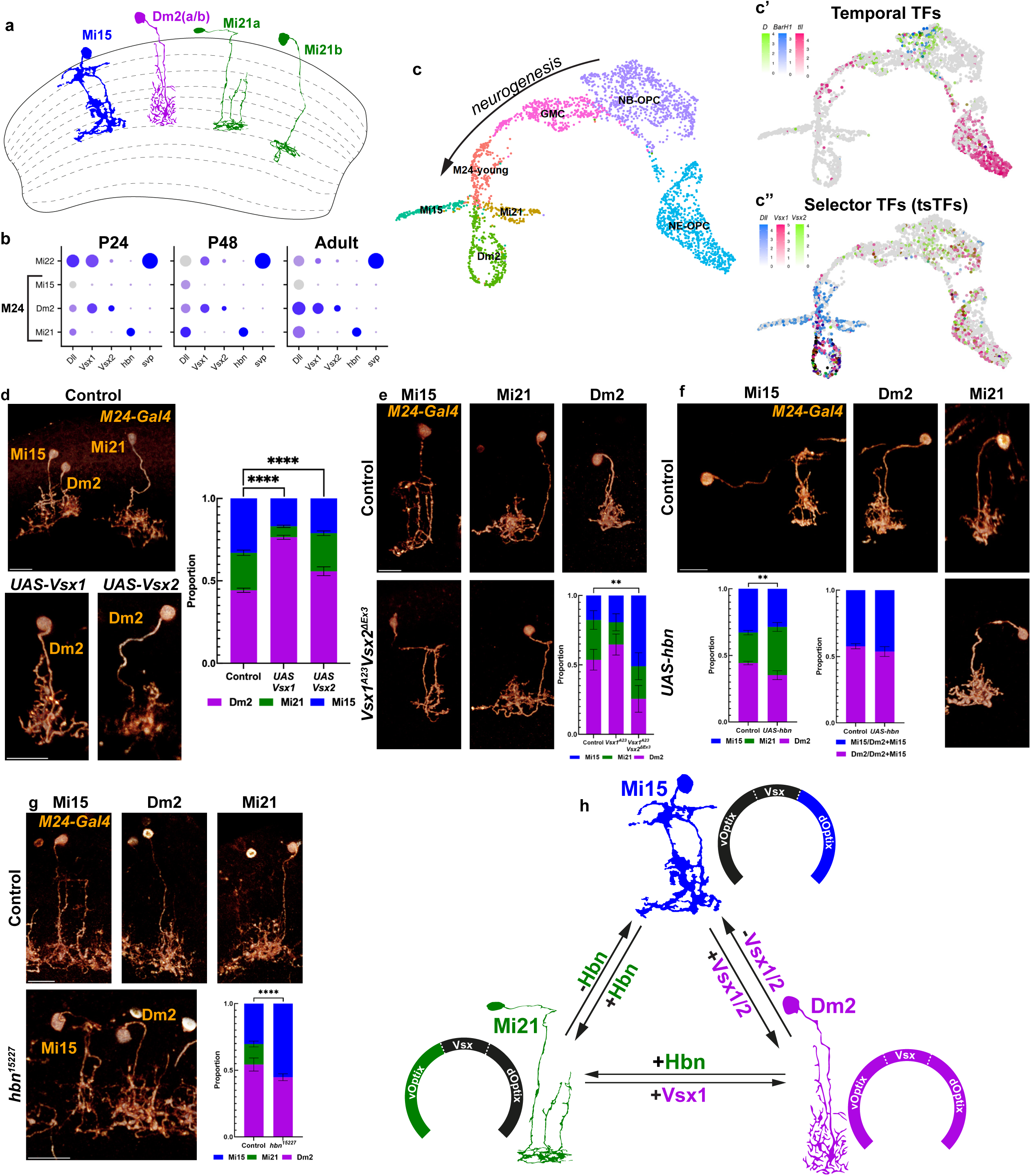
Closely related neurons Dm2, Mi15, and Mi21 can be inter-converted by terminal selectors. **a**, Cross section of the medulla neuropil showing cartoon representations of M24 cell types, Mi15 (blue), Dm2 (purple), and Mi21 (green). Of note, we recently described that Dm2 and Mi21 neurons have two very similar (a/b) subtypes resolved in our multiome atlas7. However, these subtypes do not differentially express *Vsx1/2* or *hbn*; thus, for all analyses in this study, they were considered in aggregate. Dm2 drawing was sourced from camera lucida drawings3, Mi15 from our previous study25, and Mi21a/b from FlyWire4 (Sm01 and Sm02, also known as Cm2 and Cm1). Dm2 subtypes (a/b) are morphologically indistinguishable7. **b**, mRNA expression of select marker genes for the M24 cell types (Mi15, Dm2, and Mi21) and Mi22 (also known as Cm7/Sm09) at P24, P48, and Adult stages. Dot size indicates the fraction of cells expressing the gene and color intensity indicates scaled expression level. **c**, UMAP reduction of the M24 trajectory at P0, subsetted from the NotchOFF trajectory7 and WNN calculated using 20 PCA and 24 LSI dimensions (see *Methods*), depicting shared progenitor populations of the OPC NE and NB, and a portion of the GMCs from late temporal windows, and all M24 neurons. “M24-young” cluster represents neurons that are too immature to assign identity. **c’** shows the UMAP overlaid with log-normalized mRNA expression levels of late temporal TFs *Dichaete* (*D*, green), *BarH1* (blue), and *tailless* (*tll*, pink) and **c”** selector TFs *Dll* (blue), *Vsx1* (red), and *Vsx2* (green). **d**, *M24-Gal4* driving *CD4:tdGFP* (FLP out) and *UAS-Vsx1* or *UAS-Vsx2* as annotated. n = 1108/19/12 control, n = 1997/15/9 *UAS-Vsx1*, and n = 1117/17/9 *UAS-Vsx2* neurons/optic lobes/brains. Within a quasibinomial glm, adjusted p-values for each cell type are Dm2, p<0.0001/<0.0001, Mi15, p<0.0001/<0.0001, Mi21, p<0.0001/n.s. for *UAS-Vsx1*/*UAS-Vsx2* in comparison to the control. **e**, FRT19A (control), FRT19A *Vsx1A23*, and FRT19A *Vsx1A23Vsx2ΔEx3* MARCM clones labeled with *M24-Gal4* driving *CD8:GFP* in Adult brains. Representative image not shown for the single *Vsx1A23* mutant, as it is the same as control. n = 78/23/14 control, n = 44/20/12 *Vsx1A23*, and n = 24/17/13 *Vsx1A23Vsx2ΔEx3* neurons/optic lobes/brains. Within a quasibinomial glm, adjusted p-values for each cell type are Dm2, p = n.s./ 0.00189, Mi15, p = n.s./ 0.00189, Mi21, p = n.s./n.s. for *Vsx1A23*/*Vsx1A23Vsx2ΔEx3* in comparison to the control. **f**, *M24-Gal4* driving *CD4:tdGFP* (FLP out) and *UAS-hbn* as annotated in Adult brains. Control genotype is identical to **d**. n = 1108/19/12 control and n = 686/12/6 *UAS-hbn* neurons/optic lobes/brains. Within a quasibinomial glm, adjusted p-values for each cell type are Dm2, p = 0.00214, Mi15 = n.s., Mi21, p = 0.00214 for *UAS-hbn* in comparison to the control. Second bar plot shows the relative ratio among only Dm2 and Mi15 within the M24-Gal4, ignoring Mi21. Mi15/Dm2+Mi15, p = 0.124. **g**, FRT42B (control) and FRT42B, *hbn15227* MARCM clones labeled with *M24-Gal4* driving *CD8:GFP* in Adult brains. n = 163/3 control and n = 387/6 *hbn15227* neurons/brains. Within a bias-reduced glm, adjusted p-values for each cell type are Dm2, p = n.s., Mi15, p < 0.0001, Mi21, p = 0.000632 for *hbn15227* in comparison to the control. In **d**-**g**, neurons are shown with blended three-dimensional (3D) reconstruction as representative images. All bar plots are quantifications for cell types based on marker expression and morphology with error bars representing SEM; Dm2, Mi21, and Mi15 are represented as purple, green, and blue, respectively. Scale bars: 10 μm. **h**, Cartoon representation of all conversions between M24 cell types. Beside each cartoon neuron, a spatial diagram highlights the neuroepithelial origins13 in the same color as the neuron.

We previously showed that ectopic expression of either *Vsx1* or *Vsx2* in postmitotic Mi15 neurons can morphologically and molecularly convert them into Dm2 neurons^25^; we now show that the same conversion can also be enacted in Mi21 neurons. Upon ectopic *Vsx1* expression using the *M24-Gal4* driver, we saw a drastic increase in the proportion of Dm2 neurons labeled at the expense of both Mi15 and Mi21 (**Fig. 1d**). Ectopic *Vsx2* also reduced the proportion of Mi15 (consistent with our previous results) albeit not Mi21, which is likely explained by technical reasons (**Suppl. Fig. 2**). These findings suggest that Vsx1 and Vsx2 function redundantly in specifying Dm2 identity. Consistently, Dm2 neurons were unaffected in *Vsx1* mutant MARCM (Mosaic Analysis with Repressible Cell Marker^28^) clones (**Fig. 1e**). Thus, to test this hypothesis, we generated a *Vsx2* null allele by deleting its third exon (which causes a frameshift and pre-mature stop) in the background of the existing *FRT19A*, *Vsx1^A23^* allele with CRISPR. In the MARCM clones generated using this double mutant, Dm2 were significantly depleted and Mi15 (but not Mi21) expanded (**Fig. 1e**). It is important to note that *Vsx1^A23^* does not prevent the production of Vsx1 protein but only its nuclear localization^29^, thus is likely to be a strong hypomorph rather than a null. This would explain why some Dm2 neurons could still be observed in these clones. In summary, even though ectopic *Vsx1* appears capable of converting both Mi15 and Mi21 into Dm2 morphology, upon loss of *Vsx1/2*, Dm2 neurons primarily convert to Mi15, which may be the “default” fate among these three neurons.

Next, we overexpressed the Mi21 tsTF *hbn* using *M24-Gal4*, which significantly increased the proportion of Mi21 morphology among the neurons labeled (**Fig. 1f**). This likely occurred at the expense of both Dm2 and Mi15 because their ratio was unchanged in these brains. Lastly, we generated *hbn* null MARCM clones, where Mi21 were completely eliminated with an expansion of Mi15 (but not Dm2) neurons (**Fig. 1g**). Collectively, these results establish that the expression of *Vsx1/2* in Dm2 and *hbn* in Mi21 are necessary and sufficient to differentiate these neurons from the closely related Mi15 (**Fig. 1h**). *Vsx1* could also convert Mi21 into Dm2, and vice versa with *hbn*, suggesting that these two TFs repress each other when expressed ectopically.

### Specification of neuronal Vsx1/2 expression by distinct patterning mechanisms

During neurogenesis, *Vsx1* is specifically expressed in a central domain of the OPC neuroepithelium and functions as a spatial patterning factor^14^. Accordingly, the vast majority of neurons that express *Vsx1/2* as terminal selectors are produced from the Vsx domain at L3 stage (**Fig. 2a**). Per our atlas, 38 optic lobe cell types express *Vsx1/2* as terminal selectors; lineage tracing experiments have recently demonstrated that of the 29 cell types with confidently assigned spatial origins, indeed 18 originate exclusively from this domain^13^. However, postmitotic *Vsx1/2* expression is not restricted to neurons generated from the Vsx domain and can be observed in 11 neuronal types that are produced in Optix and/or Dpp spatial domains, generally in addition to the Vsx domain^13^. Dm2 is one of these cell types that is produced from all spatial domains^15^, and consistently, they are found across the entire OPC at L3 (**Fig. 2a**, arrows).

**Figure 2:**
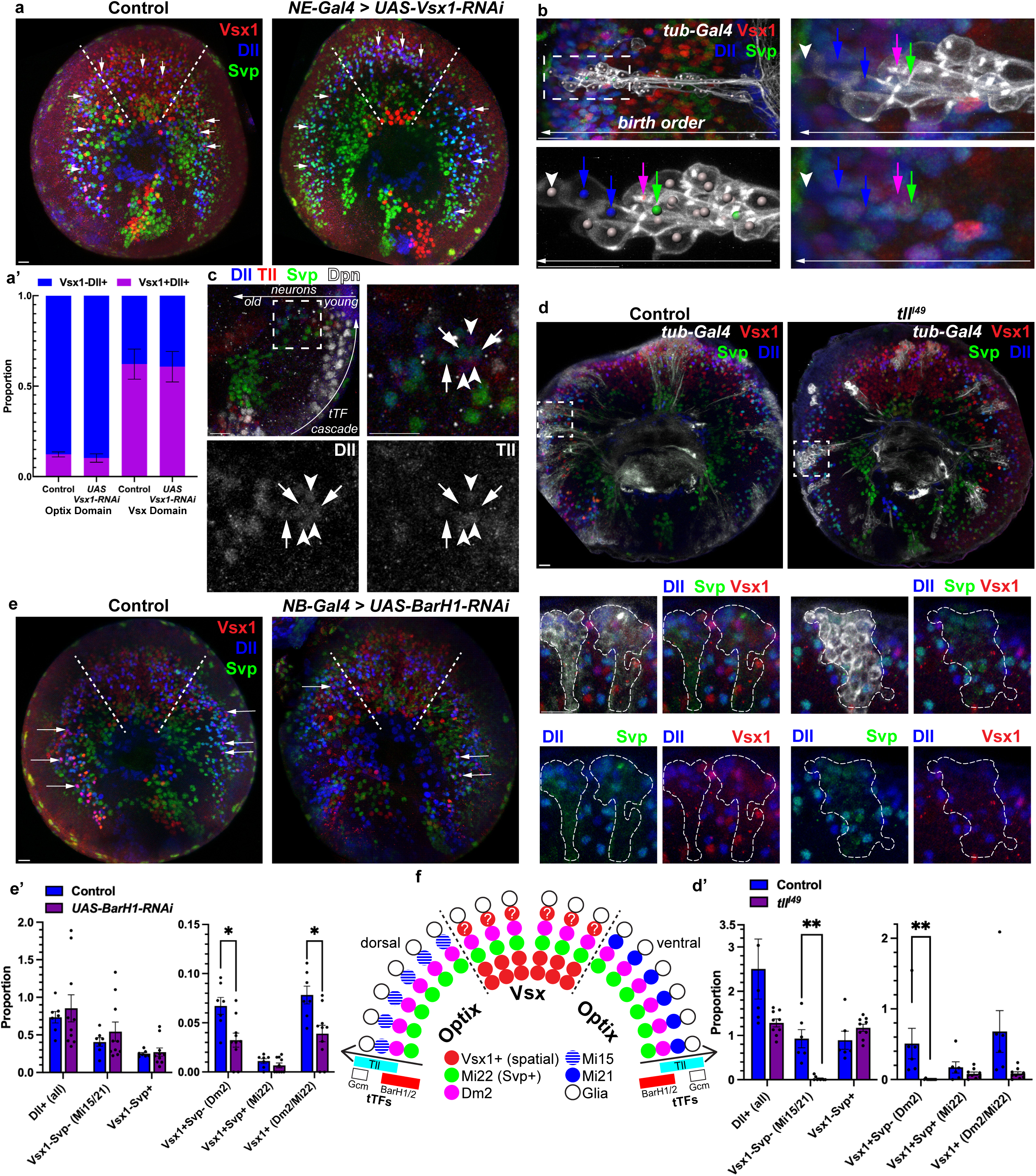
Regulation of neuronal Vsx1 expression and the birth order of Tailless window neurons a, Neuroepithelial driver (*R29C07/NE-Gal4*) driving *UAS-Vsx1-RNAi* during neurogenesis (L3) with control lacking RNAi. Dashed white lines mark the Vsx1 spatial domain and white arrows point to Dll+/Vsx1+/Svp- cells (purple, Dm2). Anti-Dll (blue), anti-Vsx1 (red), and anti-Svp (green) are shown as maximum intensity projections within a representative sample. Dm2 ratio relative to total Dll within Optix domains was not affected: 0.1227 (control) and 0.1024 (*Vsx1-RNAi*) (p = 0.473). Within the Vsx domain, average Dm2 ratio relative to total Dll was also unchanged: 0.6217 (control) and 0.6078 (*Vsx1-RNAi*) (p = 0.908). n = 154/7 control and n = 185/6 *UAS-Vsx1-RNAi* Dll+ neurons/optics lobes. Both quantifications are shown in **a’** with Dm2 shown in purple (Dll+Vsx1+) and other Dll+ neurons in blue. **b**, FRT82B MARCM clones labeled with *tub-Gal4* driving *myr:GFP* in early pupal (∼P5-P10) optic lobes. Clones are located within the dorsal Optix domain that produces all Mi15 and a portion of Dm2 neurons13. Neurons are born from NBs in sequential temporal windows from right to left marked by the white arrow. The area indicated by the white dashed box in top left panel is shown in all other panels with increased magnification. All cells marked with arrows are Dll+, the cell with green arrow is also Svp+/Vsx1+, and the cell marked with a purple arrow is Vsx1+. The arrowhead indicates glia. Anti-GFP (white), anti-Dll (blue), anti-Vsx1 (red), and anti-Svp (green) are shown as maximum intensity projections. n = 4 brains. **c**, A wild-type optic lobe at L3 shows that Tll window neuroblasts produce neurons that inherit Tll protein for a short period of time. From the bottom right to the top right, NBs (Dpn+) continuously divide and generate neurons (curved white arrow). From the NBs on the right to the left (straight white arrow), neurons are produced with age increasing along this axis. The two panels to the right show magnified views of the area indicated by the dashed white box, where arrowheads mark Dll+/Svp-/Tll+ neurons (M24) and arrows mark Dll+/Svp+/Tll- neurons. Anti-Dll (blue), anti-Tll (red or white), anti-Svp (green), and anti-Dpn (white) are shown as maximum intensity projections. n = 4 brains. **d**, FRT82B (control) and FRT82B, *tllI49* MARCM clones labeled with *tubulin-Gal4* driving *CD4:tdGFP* during neurogenesis (P5). The areas indicated in the dorsal Optix regions by white dashed boxes are shown below with higher magnification. The clones are demarcated by dashed white lines. Anti-GFP (white), anti-Dll (blue), anti-Svp (green), and anti-Vsx1 (red) are shown as maximum intensity projections. **d’**, Bar plot: Quantification of **d** with error bars representing SEM; control is blue and *tllI49* is purple. Optix domain clones were quantified relative to Otd expression in the same clones (*Methods*), with multiple clones per optic lobe quantified and summed for a single measurement per lobe. Only Dll+ cells were quantified. Adjusted p-values for each cell type are Dll+ (all), p = n.s., Dll+/Vsx1-/Svp- (Mi15/21), p = 0.00904, Dll+/Vsx1+/Svp- (Dm2), p = 0.00696, Dll+/Vsx1+/Svp+ (Mi22), p = n.s., Dll+/Vsx1+ (Dm2/Mi22), p = n.s., Dll+/Vsx1-/Svp+, p = n.s. for *tllI49* in comparison to the control. n = 6/3/23 control and n = 9/5/32 *tllI49* optic lobes/brains/clones. **e**, Neuroblast driver (*R13C02(dpn)/NB-Gal4*) driving *UAS-BarH1-RNAi* during neurogenesis (L3) with control lacking RNAi. Dashed white lines demarcate the Vsx spatial domain. Arrows point to Dm2 neurons (Dll+/Vsx1+/Svp-). Anti-Dll (blue), anti-Vsx1 (red), and anti-Svp (green) are shown as maximum intensity projections. **e’**, Quantification of **e**. ROIs within both dorsal and ventral Optix domains were quantified and normalized to Otd (*Methods*). Error bars represent SEM and each dot represents one optic lobe; control is blue and *UAS-BarH1-RNAi* is purple. Only Dll+ cells were quantified. Adjusted p-values for each cell type are Dll+ (all), p = n.s., Dll+/Vsx1-/Svp- (Mi15/21), p = n.s., Dll+/Vsx1+/Svp- (Dm2), p = 0.0257, Dll+/Vsx1+/Svp+ (Mi22), p = n.s., Dll+/Vsx1+ (Dm2/Mi22), p = 0.0257, Dll+/Vsx1-/Svp+, p = n.s. for *UAS-BarH1-RNAi* in comparison to the control. n = 13/7/6 control and n = 15/10/9 *UAS-BarH1-RNAi* Optix domains/ lobes/ biological replicates. Scale bars: 10 μm. We presume that *Vsx2* expression aligns with *Vsx1* in these experiments because these genes are always co-expressed in optic lobe neurons25, but we did not explicitly test this. **f**, Graphical representation of M24 cell types in the context of spatial and temporal patterning of the OPC. The postmitotic neurons (circles) extend in concentric half-circles representative of a NB producing progeny along a temporal axis (arrows) with spatial domains annotated in bold and delineated by dashed lines. Along the temporal axis, boxes represent the relevant temporal TFs’ expression and their overlap: BarH1/2 (red), Tll (cyan), and Gcm (white). Red circles represent Vsx1 expressing progeny that arise exclusively from the Vsx spatial domain. Green circles represent Mi22 neurons which express Dll, Svp, and Vsx1 and are born before M24 neurons. Purple circles represent Dm2 which are born during the BarH1/2 and Tll overlap. Blue circles represent the latest born M24 neurons, Mi15 and Mi21. Mi15 (dorsal Optix) is shown with horizontal stripes while Mi21 neurons (ventral Optix) are predicted to be produced during the same temporal window. Red circles with white question marks depict the unknown cell type that may be produced from the Vsx domain during this temporal window. White circles depict the glia produced last during the overlap of Tll and Gcm17.

The strong correlation with domain origin and Vsx1/2 expression led to the attractive hypothesis that postmitotic Vsx expression is instructed by neuroepithelial Vsx1. *Vsx1* was indeed shown to be necessary for specification of Pm3^14^, which is one of these domain-exclusive neurons. However, the previous experimental designs could not separate neuronal vs. neuroepithelial roles, as the knockdowns were permanent in the lineages they were induced. Thus, we performed a transient, NE-specific knockdown of *Vsx1*, which drastically reduced Vsx1 in neurons emerging from its spatial domain, including the loss of all early-born neurons that co-express Vsx1 and TfAP-2 (Pm3, TmY15, TmY17, and Tm23) (**Suppl. Fig 3a**). This result confirms that Vsx1-mediated patterning of the OPC neuroepithelium indeed instructs *Vsx1/2* expression in many neurons that originate from the corresponding spatial domain. However, this is not due to a simple inheritance or maintenance of its expression (which we further discuss below), as Vsx1 is downregulated in NBs before being re-activated in only some of the neurons that originate from the Vsx domain^14,29^. More importantly for this study, NE knockdown did not affect *Vsx1* expression in Dm2 neurons, including those within the Vsx domain (**Fig. 2a**, arrows). Therefore, *Vsx1* expression in Dm2 neurons must be regulated independently from Vsx1-mediated spatial patterning.

While the Vsx1/2^+^ Dm2 neurons are produced from all spatial domains, the related Vsx^-^ cell types Mi15 and Mi21 (discussed above) originate from mutually exclusive Optix domains (Mi15: dorsal, Mi21: ventral^13^, **Fig. 1h**). As all M24 neurons have the same N^OFF^ status, we hypothesized that differential *Vsx1/2* expression among them is instructed by temporal TFs. To understand this regulation, we first sought to resolve the birth order of these neurons by analyzing sparse NB clones in the dorsal Optix domain (**Fig. 2b**, **Suppl. Fig. 3b**). Dll is a tsTF expressed specifically in N^OFF^ OPC neurons produced near the end of the temporal cascade^12^ and in all spatial domains, including all M24 neurons (**Fig. 1b-c**). Among the cell types that are produced in dorsal Optix^13^ and express *Dll*, we observed that Mi22 neurons^7^ that co-express Vsx1 and Svp (but are not part of M24, **Fig. 1b**) are produced first (**Fig. 2b**, green arrow), followed by Dm2 that express Vsx1 (but not Svp, **Fig. 2b**, purple arrow) and then Mi15 (Dll only, **Fig. 2b**, blue arrows). The glia that are produced during the final temporal window of OPC neuroblasts^18^ (**Fig. 2b**, arrowhead) also express *Dll*, but the protein is not detectable before their migration towards the neuropil immediately after their birth (**Suppl. Fig. 3c**). Consistent with these results, all M24 neurons (Dll^+^/Svp^-^), but none of the Svp^+^ neurons, transiently express the late temporal TF Tailless (Tll) protein inherited from NBs (**Fig. 2c**), similar to the glia produced from Tll^+^ NBs^18^. Further supporting this, in *tll* mutant MARCM clones, the only neurons that still expressed Dll also expressed Svp (**Fig. 2d**), indicating the loss of all M24 neurons. The glial cells (Repo^+^) were similarly lost in these mutant clones (**Suppl. Fig. 3d**). Because *tll* is required to produce all M24 neurons, we concluded that differential *Vsx1/2* expression in Dm2 compared to Mi15 and Mi21 must be regulated by another temporal factor.

The expression of temporal TFs BarH1/2 precedes and partially overlaps with Tll expression in NBs^12,17^. Accordingly, *BarH1* RNAi knockdown in NBs significantly reduced the number of Dm2 neurons produced (**Fig. 2e**). In contrast, the number of Mi15 and Mi21 (Dll^+^/Vsx1^-^/Svp^-^) neurons was slightly (but not significantly) increased (**Fig. 2e’**). Upon *BarH1* and *BarH2* double knockdown in NBs, all Dll^+^ (including M24) neurons were significantly reduced (**Suppl Fig. 3e**). This is likely an indirect effect consistent with previous observations that BarH1/2 are redundantly necessary also for *tll* activation in NBs^12,17^. These results indicate that BarH expression during the first part of the Tll window directly or indirectly activates *Vsx1* expression in Dm2, distinguishing them from Mi15 and Mi21. Together, these results show that the OPC NBs expressing *tll* have at least three temporal windows (defined by the overlapping expression of *BarH* or *gcm*^17^) for sequential production of first Dm2, then Mi15 (and presumably Mi21) and then finally glia (**Fig. 2f**).

### Modular regulation of neuronal Vsx1/2 expression by distinct enhancers

We next investigated the cis-regulatory mechanisms of neuronal *Vsx1/2* expression. Using our single-cell multiome atlas of the developing optic lobes^7^, we analyzed expression and accessibility of the *Vsx1/2* locus at P0 during neurogenesis. We observed that a previously described domain-specific enhancer of this locus^30^ was specifically accessible in the Vsx domain NE, but not accessible in NBs or GMCs (**Fig. 3a**, brown rectangle). Despite having some accessibility also in the Vsx domain neurons, a concurrent study by Chen and colleagues^31^ has demonstrated that the activity of this enhancer is indeed restricted to the neuroepithelium. Notably, two other regions that display domain-specific accessibility in the NE were completely closed in neurons (**Fig. 3a**, purple). However, we found multiple (other) regions that were consistently accessible in the Vsx^+^ neurons that originate from the Vsx domain and highlighted two of them (**Fig. 3a**, blue); these, in contrast, were largely not accessible in OPC progenitors. The neuron-specific activity of these enhancers were also recently validated with *in vivo* reporters^31^. These extend our recent findings on temporal TFs^7^ to the spatial TFs: even when these progenitor patterning factors appear to be “maintained” as terminal selectors in the corresponding neuronal progeny, they are rather “re-activated” through separate enhancers.

**Figure 3:**
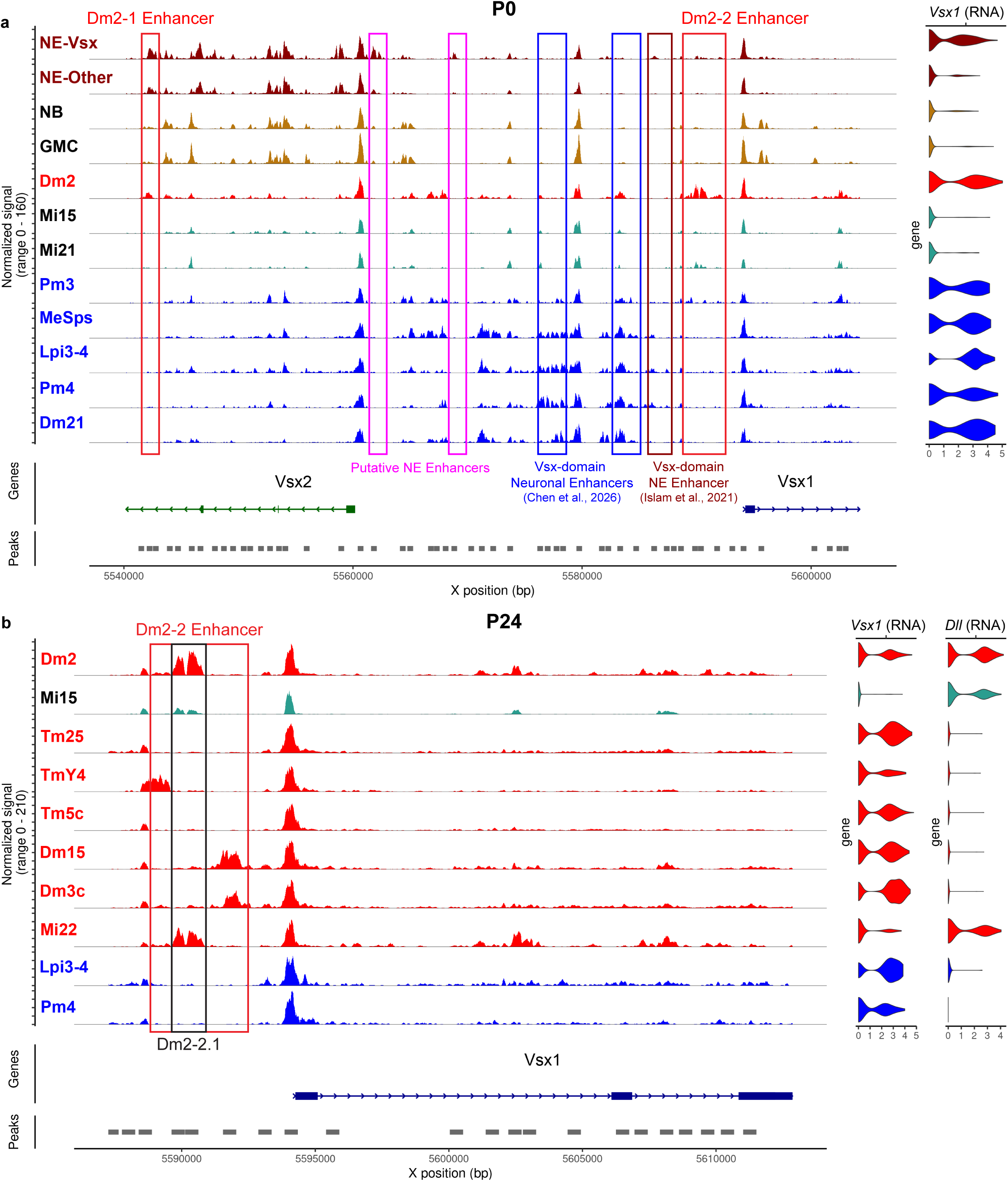
Accessibility of temporally patterned enhancers of the *Vsx* locus. **a**, Coverage plot of the *Vsx1/2* locus with tracks representing normalized accessibility in each cell type at P0. Neuroepithelium (NE, dark red) has been split into Vsx1+ and other domains. NB: neuroblast; GMC: ganglion mother cell. Five representative Vsx1+ cell types that exclusively originate from the Vsx1 spatial domain are shown in blue (two Pm3 subtypes were merged). Rectangles highlight Dm2-1 and Dm2-2 enhancers with Dm2-specific accessibility (red) and the enhancers commonly accessible within spatially defined cell types (blue) as described in Chen et al.31. NE enhancers, including the previously described Vsx-domain enhancer30 (dark red) and other putative NE enhancers (purple) are also shown. Log-normalized *Vsx1* mRNA expression levels in each cell type are shown on the right side.**b**, Coverage plots of the P24 multiome data show normalized accessibility of the ∼4kb Dm2-2 enhancer region in Dm2 and Mi15 neurons and additional *Vsx1/2*-expressing cell-types. By P24, accessibility becomes restricted to distinct ∼1 kb subregions specific to each lineage; Dm2-2.1 (black) is the subregion for BarH temporal window (Dm2/Mi22). Violin plots (right) display log-normalized mRNA expression of *Vsx1* and *Dll* in these clusters. Lpi3-4 and Pm4 (blue) are representative cell types that exclusively originate from the Vsx domain and are thus spatially patterned while cell types in red originate from both Vsx and Optix domains. We note that there are some neurons (e.g. Tm5c and Tm25) that express *Vsx1/2* despite originating from other domains, but do not appear to use this enhancer cluster. This suggests that there are likely additional enhancers of *Vsx1/2* that are temporally patterned.

Our findings in the previous section predict that *Vsx1/2* expression in Dm2 should be regulated differently from the Vsx-domain neurons. Among the M24 neurons, we identified two regions in the *Vsx1/2* locus that are highly accessible in Dm2 (from P0 to Adult) and with low or no accessibility in Mi15, Mi21, or in any of the Vsx^+^ neurons that originate exclusively from the Vsx domain (**Fig. 3a**, red). We generated *Gal4* reporters for both enhancers and analyzed their expression patterns throughout optic lobe development (**Fig. 4, Suppl. Fig. 4**). The reporter for the first enhancer (Dm2-1, within the 3^rd^ intron of *Vsx2*) stochastically labeled a minority of Dm2 neurons (on average 17% in Adults) from P24 to Adult and very few OPC neurons at the L3 stage during neurogenesis (**Fig. 4a** and **Suppl. Fig. 4a,c**). The reporter for the second enhancer (Dm2-2, ∼4kb upstream of *Vsx1*) labeled Dm2 neurons much more consistently throughout development (on average 48% in Adults), including at L3 (**Fig. 4b-d** and **Suppl. Fig. 4b,d**). The overall expression pattern of the Dm2-2 reporter was concentric (i.e. spanning all spatial domains) during neurogenesis (**Fig. 4b’**), consistent with temporal patterning of *Vsx1/2* expression in Dm2 neurons. Both reporters also labeled neurons beyond Dm2, including some that lacked Vsx1; we discuss the origins of this off-target activity in **Supplementary Note 1**.

**Figure 4:**
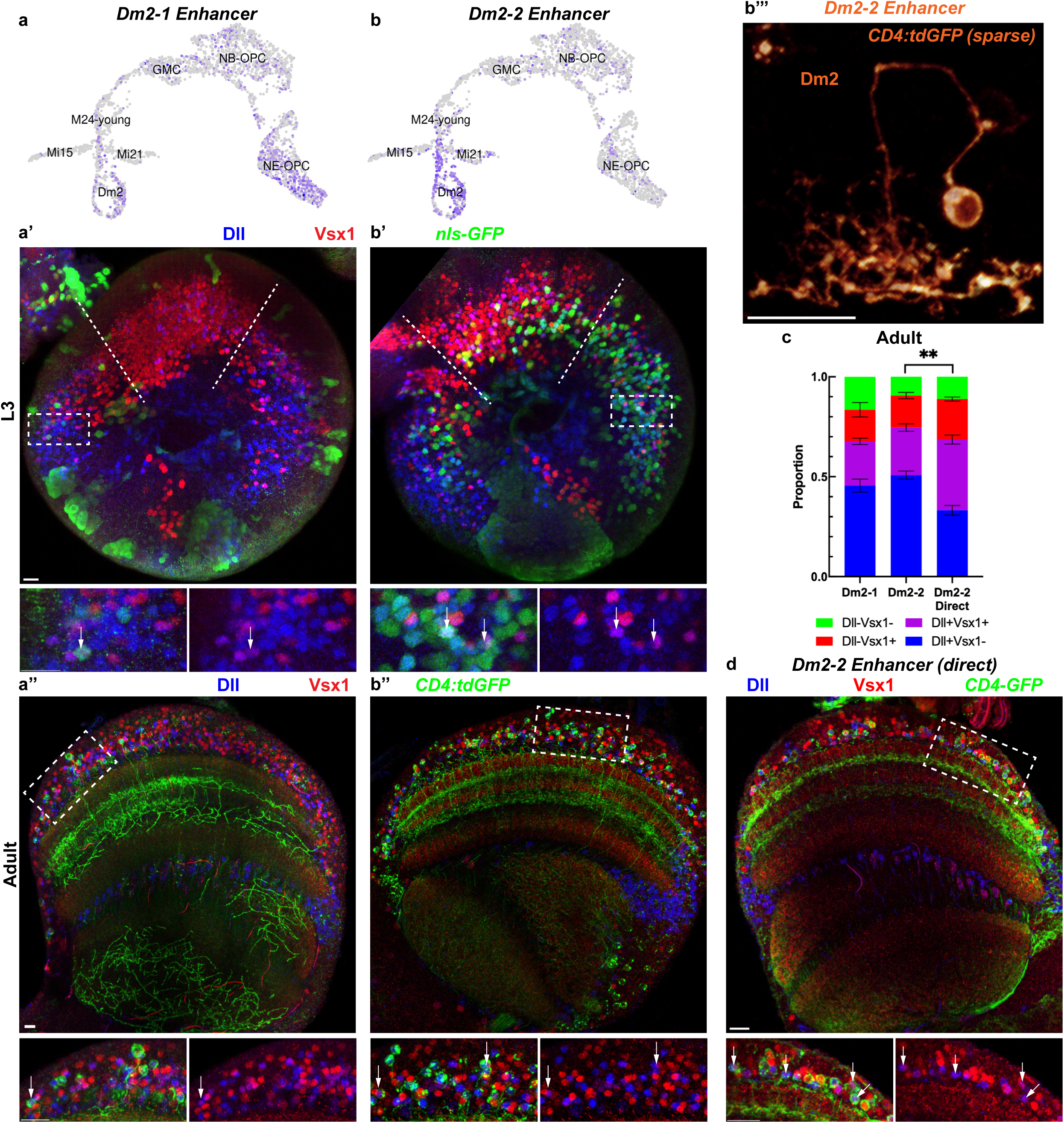
Activity of temporally patterned enhancers of the *Vsx* locus. **a-b**, Dm2-1 and Dm2-2 enhancer accessibility displayed on the same UMAP as in Fig. 1c with cell types annotated. “M24-young” refers to neurons too immature to confidently assign identity. Both enhancers were cloned into a *Gal4* reporter driving nuclear (nls) GFP within L3 brains (**a’** and **b’**, Dm2-1, n = 5 and Dm2-2, n = 5 brains) and membrane-tagged (CD4) GFP within adult brains (**a”** and **b”**, Dm2-1, n = 5 and Dm2-2, n = 6 brains). Representative images from each stage are shown as maximum intensity projections with anti-GFP (green), anti-Dll (blue), and anti-Vsx1 (red). White arrows point to Dll+/Vsx1+ co-expression and white dashed lines in L3 brains demarcate the Vsx1 spatial domain. See also **Suppl. Fig. 4** for all reporters in all stages (L3-Adult). **b”’**, 3D reconstruction of an adult Dm2 neuron sparsely labeled with *Dm2-2-Gal4* driving (FLP-out) *CD4:tdGFP*. n = 8 brains. **c**, Quantification of adult brains from **a**,**b**, and **d** for Dm2-1 and Dm2-2 (*Gal4* and direct) enhancers was performed as a relative proportion of all cells labeled by the reporter (*Methods*) with error bars indicating SEM. Dll-/Vsx1-, Dll-/Vsx1+, Dll+/Vsx1+, and Dll+/Vsx1- cells are shown in green, red, purple, and blue, respectively. n = 516/5 (Dm2-1), n = 816/6 (Dm2-2), and n = 952/4 (Dm2-2-direct) neurons/brains. Quasibinomial glm comparison of Dm2-1 and Dm2-2 returned no significant values; Dm2-2 (*Gal4*) and Dm2-2-direct adjusted p-values Dll-/Vsx1-, p = n.s., Dll-/Vsx1+, p = n.s., Dll+/Vsx1+, p = 0.00418, and Dll+/Vsx1-, p = 0.00294. **d**, Dm2-2 enhancer was cloned into a reporter directly driving expression of membranous GFP and visualized within Adult brains (n = 4). Anti-Dll (blue), anti-Vsx1 (red), and anti-GFP (green) are shown as maximum intensity projections. Dashed white region is shown with greater magnification to the right with white arrows pointing to Dll+/Vsx1+ cells (Dm2). Scale bars: 10 μm.

The majority of the cells labeled by these reporters in the medulla expressed Dll (**Fig. 4c**). Since Dll in the OPC is highly specific to the N^OFF^ neurons from late temporal windows^12^, this result further highlights the temporal bias in these enhancers’ activity. However, both Dm2-1 and Dm2-2 reporters were also expressed in some Vsx1^+^ neurons that were Dll^-^ (i.e. not Dm2), which prompted us to investigate the accessibility of these enhancers more generally in the OPC. We focused on the Dm2-2 enhancer because its activity was higher than Dm2-1 and appeared to commence earlier (**Fig. 4a-b**), thus it is more likely to control the initiation of *Vsx1/2* expression. ATAC signal in this enhancer was strong in at least four other neurons in addition to Dm2: *Dll*^+^ Mi22, and *Dll*^-^ TmY4, Dm15, and Dm3c (**Fig. 3b, Suppl. Fig. 5**), all of which express *Vsx1/2* despite also being produced in both Vsx and Optix spatial domains^13^. In stark contrast, Dm2-2 was not accessible in *any* of the neurons that are produced exclusively in the Vsx domain (**Fig. 3b**, **Suppl. Fig. 5**, Pm4 and Lpi3-4 are representative examples).

Interestingly, while this ∼4kb region was broadly open in Dm2 at P0 (**Fig. 3a**, **Suppl. Fig. 5**), the accessibility became restricted to a ∼1kb peak within this region from P24 onwards (**Fig. 3b**, **Suppl. Fig. 5**). Similarly, in TmY4 (N^ON^, Slp/D temporal window), it became restricted to a distinct peak upstream of the Dm2 peak; Dm15 and Dm3c (N^OFF^, Slp/D) shared a different peak downstream of the Dm2 peak (all within the same 4kb region). N^OFF^ Mi22 neurons shared the same subregion as Dm2. We showed above that Mi22 neurons are produced immediately before Dm2, thus they are likely also born from BarH-expressing NBs (but before Tll expression, **Fig. 2b**). While the reduction in the number of these neurons upon *BarH1* knockdown was not significant (**Fig. 2e’**), we note the rarity of this cell type (only ∼90 cells per optic lobe, compared to ∼800 Dm2 or Mi15+Mi21^5^), resulting in infrequent representation and lower statistical power. Therefore, these findings are consistent with the hypothesis that BarH1 specifies *Vsx1/2* expression in both Dm2 and Mi22, but it could also reflect their shared expression of Dll. We generated a *Gal4* reporter for this specific sub-region of the Dm2-2 enhancer (named Dm2-2.1, **Fig. 5c-d**). In adult brains, the proportion of Dm2/Mi22 (Dll^+^/Vsx1^+^) neurons labeled was dramatically higher compared to the full Dm2-2 reporter (**Figs. 4c**, **5d’**), and indeed 95% of all neurons labeled by the reporter expressed Dll (**Fig. 5d’**). At L3, in contrast to the full Dm2-2 reporter (**Fig. 4b**), Dm2-2.1 activity was strikingly depleted from the Vsx spatial domain, but it still labeled Dm2 neurons in the Optix domain (**Fig. 5c**).

**Figure 5:**
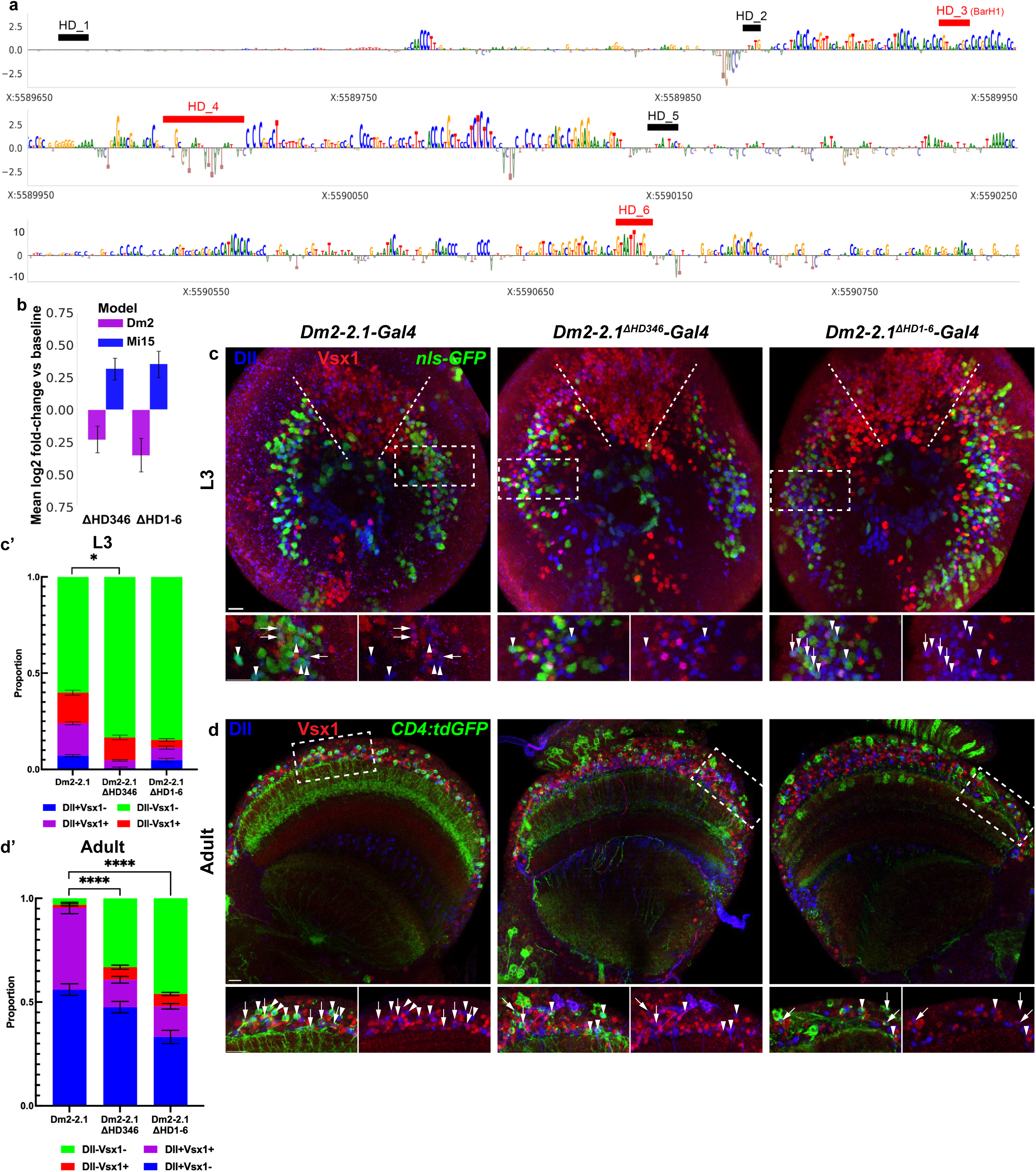
Deep learning guided perturbation of TF binding sites. **a**, Base-pair contribution scores (deepSHAP36) within the *Dm2-2.1* enhancer per the P0 Dm2 ChromBPNet model, with the positions of six potential HD motifs (per FIMO ran on reference dm6, see Suppl. Table 1) indicated. Those in red represent the three highest-importance motifs per the ChromBPNet models and deleted specifically in the *Dm2-2.1ΔHD346* reporter (c). **b**, Predicted accessibility changes (compared to the wild-type sequence) for the *Dm2-2.*1 enhancer upon deletion of either motifs 3,4,6 or all (1-6) HD motifs annotated in (**a**), in Dm2 and Mi15 neurons at P0 per the corresponding ChromBPNet models. The average predicted accessibility changes over five independent models per cell type is shown with error bars representing standard deviation. **c**-**d**, *Dm2-2.1-Gal4*, *Dm2-2.1ΔHD346-Gal4*, and *Dm2-2.1ΔHD1-6-Gal4* driving nuclear (nls) GFP within L3 brains (Dm2-2.1 = 6, ΔHD346 = 4, ΔHD1-6 = 5 brains) and membrane tagged (CD4) GFP within adult brains (Dm2-2.1 = 6, ΔHD346 = 5, ΔHD1-6 = 5 brains). Representative images from each stage are shown as maximum intensity projections with anti-GFP (green), anti-Dll (blue), and anti-Vsx1 (red). White arrows point to Dll+/Vsx1+ co-expression and white arrowheads point to Dll+/Vsx1- expression. Scale bars: 10 μm. Quantification of cells labeled at L3 (**c’**) and adult (**d’**) was performed as a relative proportion of all cells labeled by the reporter (*Methods*) with error bars indicating SEM. Dll-/Vsx1-, Dll-/Vsx1+, Dll+/Vsx1+, and Dll+/Vsx1- cells are shown in green, red, purple, and blue, respectively. For Adult, adjusted p-values are Dll+/Vsx1-, p = n.s./0.000141/0.0088, Dll+/Vsx1+, p = <0.0001/<0.0001/n.s., Dll-/Vsx1+, p = 0.00207/0.00145/n.s., Dll-/Vsx1-, p = <0.0001/<0.0001/0.0179 for Dm2-2.1 vs ΔHD346/ Dm2-2.1 vs ΔHD1-6/ΔHD346 vs ΔHD1-6. For L3, adjusted p-values are Dll+/Vsx1-, p = 0.00447/0.0211/n.s., Dll+/Vsx1+, p = 0.0220/n.s./n.s., Dll-/Vsx1+, p = n.s./n.s./n.s., Dll-/Vsx1-, p = 0.0183/0.0211/n.s. for Dm2-2.1 vs ΔHD346 / Dm2-2.1 vs ΔHD1-6 / ΔHD346 vs ΔHD1-6.

Together, these findings indicate that different enhancers have evolved, in curiously close proximity within the Dm2-2 region, to activate *Vsx1/2* expression in neurons from different temporal windows (and Notch status), and with complete independence from Vsx1-mediated patterning of the neuroepithelium.

### Deep-learning guided validation of key binding sites within the temporal *Vsx1/2* enhancer

Our multiome dataset^7^ enabled us to identify the enhancers that likely regulate *Vsx1/2* in optic lobe neurons. However, it remains unclear whether the temporal TF BarH1 acts directly through the Dm2-2 enhancer for the activation of *Vsx1/2* expression in Dm2 neurons. Furthermore, considering that BarH1 (like most temporal TFs) is not maintained in differentiated neurons (**Fig. 1c’**), it is also unclear how the accessibility and activity of this enhancer is maintained until adulthood. Notably, BarH1/2, Vsx1/2 and Dll are all homeodomain (HD) TFs with highly similar consensus binding motifs. Motif scanning (using FIMO^32^) for known *Drosophila* PWMs revealed at least 6 different potential HD binding sites within the *Dm2-2.1* region (**Suppl. Table 1**), including one (HD_3) direct match to the BarH1 SELEX PWM^33^. Three of these (HD_1/4/6) were composite sites, containing multiple copies of the core TAAT/ATTA motif, which could enable homo- or heterodimer binding as typical of HD TFs^34^.

Because most genomic instances of TF motifs are not necessarily functional binding sites, we trained ChromBPNet^35^ models on the chromatin accessibility profiles of Dm2 and Mi15 neurons at P0. These convolutional deep learning algorithms learn to predict cell-type-specific accessibility from DNA sequence. Once trained, they can accurately predict ATAC signals of even held-out sequences (see *Methods*), and they can be interrogated to reveal the specific base-pairs in each enhancer with the highest contributions towards model predictions, often representing important TF binding sites. Three of the 6 potential HD binding sites (HD_3/4/6) received high (positive or negative) deepSHAP^36^ contribution scores from the P0 Dm2 ChromBPNet model (**Fig. 5a**). We regenerated the *Dm2-2.1-Gal4* reporter with these motifs deleted (*Dm2-2.1^ΔHD346^*). As predicted by the models (**Fig. 5b**), the proportion of Dm2 (Dll^+^/Vsx1^+^) neurons labeled by the reporter was strongly and specifically reduced in the deletion construct compared to wildtype in both L3 (**Fig. 5c**) and adult (**Fig. 5d**) optic lobes. As a control, we also deleted all potential HD sites identified by FIMO in this enhancer (*Dm2-2.1^ΔHD1-6^*). We did not observe any additional depletion of Dm2s labeled by this reporter compared to *Dm2-2.1^ΔHD346^*-*Gal4* at L3 or Adult (**Fig. 5c’, d’**), indicating that ChromBPNet models were indeed able to decipher the key binding sites in the Dm2-2 enhancer.

As there are no other HD TFs differentially expressed between Dm2 and Mi15, we conclude that after the initial activation by BarH1, Vsx1/2 can maintain their own expression through the Dm2-2 enhancer. However, our results (i.e. the restricted activity of *Dm2-2.1-Gal4* in the late-born N^OFF^ neurons) also indicate that this auto-regulation would have to be context dependent and likely requires the cooperative action of Vsx1/2 and the shared M24 tsTF Dll.

### Spatial regulation of a temporal transcription factor

Both the previous lineage tracing experiments^13^ and our results above support that N^OFF^ Mi15 and Mi21 neurons, which are distinguished by the tsTF Hbn (**Figs. 1b,g**, **6b**), are produced in dorsal and ventral Optix spatial domains, respectively (**Fig. 6a**), within the same temporal window (**Fig. 2f**). Hbn is also a temporal TF of OPC neuroblasts, but as we previously described^7,12^, it is not expressed in any of the neurons that originate from the Hbn temporal windows, and all neurons that express it as a tsTF (including Mi21) are born during the later temporal windows. Thus, while the neuroblast vs. neuronal regulation is similarly expected to be independent for *hbn*, our findings raise the interesting possibility that, just as the spatial TF *Vsx1* is regulated by temporal factors in some neurons, the temporal TF *hbn* could be regulated by spatial patterning in Mi21.

**Figure 6:**
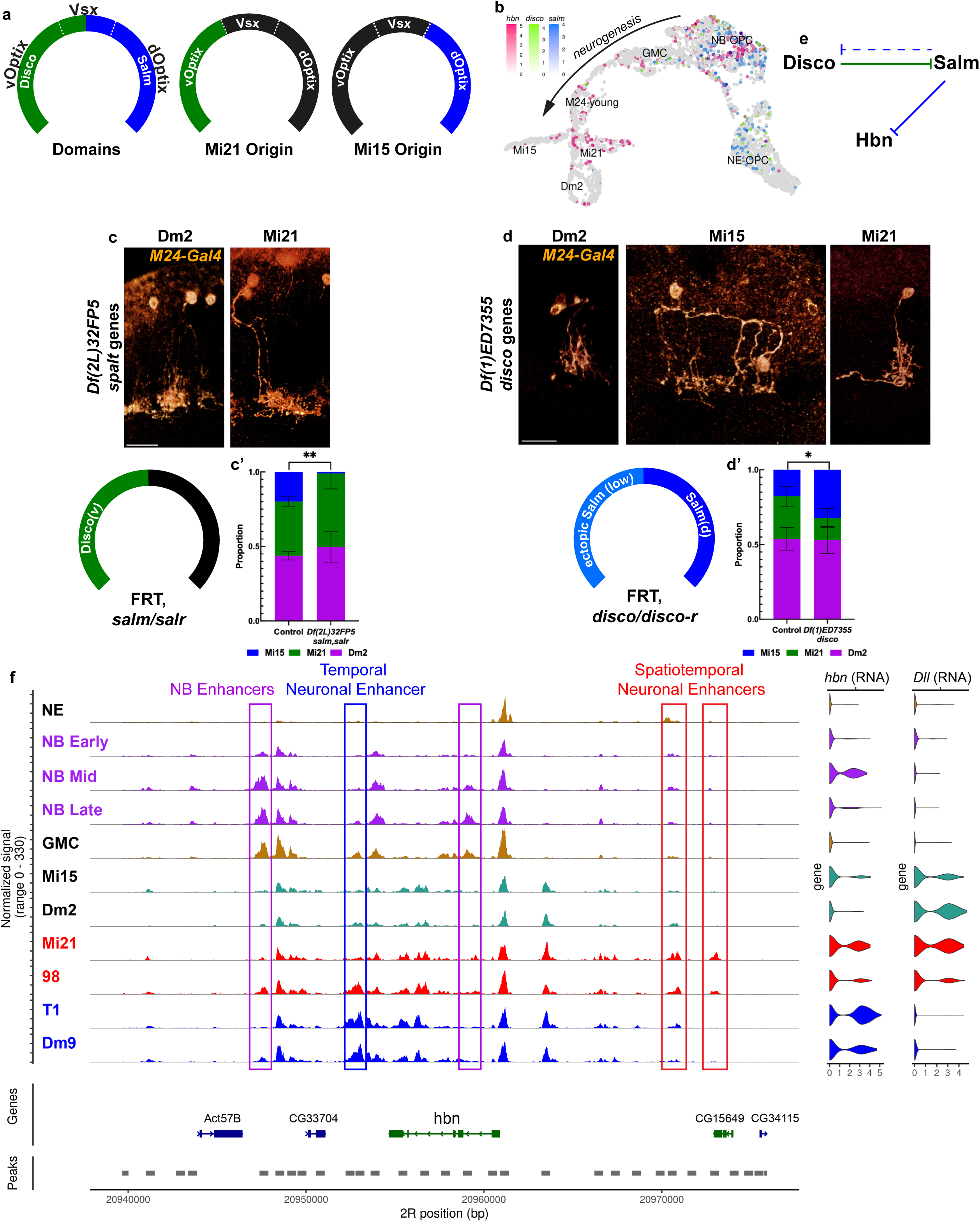
Spatial regulation of the temporal transcription factor Hbn. **a**, Spatial patterning of the OPC neuroepithelium. White dashed lines delineate Vsx and Optix domains while green and blue represent dorsoventral patterning by Disco and Salm. Mi15 and Mi21 spatial origins are replicated from Fig. 1h. **b**, Expression of tsTF *hbn* (red) and neuroepithelial TFs *disco* (green) and *salm* (blue) on the same UMAP as in Fig. 1c with cell types annotated. In **c**-**d**, cell types are shown in anti-GFP (orange) with blended 3D reconstruction and annotated as a representative image. Diagrams depict *disco* and *spalt* genes expression change in each experimental condition as described in a previous study16. **c**, FRT40A (control) and FRT40A, *Df(2L)32FP5* (covering *salm*/*salr*) MARCM clones labeled with *M24-Gal4* driving *CD8:GFP* in adult brains. **c’,** Bar plot: quantification of **c** with error bars representing SEM. In comparison to the control, adjusted p-values are Dm2, p = n.s., Mi15 = 0.0025, Mi21, p = n.s. for *Df(2L)32FP5.* n = 841/10/5 control and n = 298/8/5 *Df(2L)32FP5* neurons/optic lobes/brains. **d**, FRT19A (control) and FRT19A, *Df(1)ED7355* (covering *disco*/*disco-r*) MARCM clones labeled with *M24-Gal4* driving *CD8:GFP* in adult brains. **d’,** Bar plot: quantification of **d** with error bars representing SEM. In comparison to the control, adjusted p-values are Dm2, p = n.s., Mi15, p = 0.0263, Mi21, p = n.s. for *Df(1)ED7355*. n = 78/23/14 control and n = 111/9/5 *Df(1)ED7355* neurons/optic lobes/brains. Control genotype is identical to Fig. 1e. Scale bars: 10 μm. **e**, Proposed regulation of Mi21 Hbn expression. Disco and Salm co-repress each other as shown in a previous study16 and Salm loss does not lead to Disco expansion (dashed line). **f**, Coverage plot of the *hbn* locus with tracks representing normalized accessibility in each cell type at P0. All neuroblast subclusters (Early, Mid, Late) are shown in purple. NE, GMC, Dm2 and Mi15, which do not express *hbn*, are written in black. All other cell types express *hbn* as a tsTF. Mi21 and cluster 98 (red) are later born cell types from vOptix domain. T1 and Dm9 (blue) are produced from all spatial domains. Rectangles highlight putative cis-regulatory regions for NBs (purple), domain-agnostic neurons (blue), and spatially-patterned neurons (red). Log-normalized *hbn* and *Dll* mRNA expression levels in each cell type are shown on the right side.

*Salm/salr* (*spalt* genes) and *disco/disco-r* (*disco* genes) divide the OPC neuroepithelium into its dorsal and ventral (d/v) regions, respectively^16^ (**Fig. 6a**). These domains overlay Optix and Vsx1 regionalization, adding further spatial inputs for cell-type patterning. However, unlike *Vsx1*, *spalt* and *disco* genes are never expressed postmitotically in OPC neurons (**Figs. 1c**, **6b**). To investigate *spalt* and *disco* genes as potential spatial regulators of *hbn* among temporally similar M24 neurons, we induced null MARCM clones using *M24-Gal4* and quantified the abundance of each cell type in adult brains. First, we observed that loss of *spalt* eliminated almost all Mi15 neurons that originate from dOptix (**Fig. 6c**). Importantly, in these brains the Mi21:Dm2 ratio among the labeled neurons was higher (**Fig. 6c’**), suggesting that upon loss of *spalt* Mi15 neurons were not simply lost but indeed converted to Mi21 fate. Next, we analyzed the effects of *disco* loss using the same approach and found that Mi15 proportion was significantly increased in the *disco* mutant clones (**Fig. 6d’**). Importantly, it was previously shown that while the loss of *disco* de-represses *salm* and expands its expression to the ventral NE (albeit at lower levels than the dorsal side), loss of *spalt* does not de-repress *disco* in the dorsal NE (it remains restricted to ventral half)^16^. This partial de-repression of *salm* upon *disco* loss likely explains the finding that Mi21 were still present in that condition. Together, these results are consistent with a model whereby within the Tll temporal window, *spalt* is responsible for repression of *hbn* in the dOptix domain, distinguishing Mi15 neurons from their ventral counterpart Mi21 (**Fig. 6e**).

Within the *hbn* locus, there are several candidate enhancers that could regulate its expression in both the NBs and postmitotic neurons. As a temporal TF, *hbn* is expressed in the “middle” part of the temporal cascade and there are multiple enhancers whose accessibility is restricted to the middle and late group of NBs, likely responsible for its regulation as a temporal TF (**Fig. 6f**). Mi21 and an unannotated neuron type (cluster 98 in our multiome atlas^7^) are produced during the late temporal windows from the vOptix domain. Cluster 98 expresses *Dll* and *Svp*, implying its similarity to Mi21 neurons and slightly earlier birth before the Tll window (**Fig. 2d**). Both Mi21 and cluster 98 share accessibility in some *hbn* enhancers that are not open in progenitors (**Fig. 6f**). These enhancers are also not accessible in other Hbn^+^ cell types that are produced from all spatial domains (e.g. T1 and Dm9); those cell types seem to use separate enhancers to regulate their expression of *hbn* as a tsTF.

Taken together with our investigation of *Vsx1/2* regulation, we have found not a rare exception, but a pattern that tsTF expression in postmitotic neurons is highly complex and controlled by diverging cis-regulatory elements. This allows their independent regulation at different developmental steps (progenitor vs. neuron) and, in neurons, downstream of different patterning mechanisms (spatial vs. temporal).

## Discussion

The *Drosophila* optic lobe is a uniquely powerful model system to understand how terminal selector combinations are specified: the spatial, temporal and Notch origins of most OPC neurons have now been mapped, their putative tsTF codes inferred^25^, and the correlations between developmental origin and selector expression established^13^. However, the cis-regulatory logic that converts a neuron’s developmental origin into the activation of the correct set of terminal selectors is only starting to become understood. By dissecting the regulation of a single locus, we show here that this conversion does not follow a simple mapping: *Vsx1/2* are activated as terminal selectors by spatial patterning in most OPC neurons but by the temporal TF BarH1 in Dm2, through physically distinct enhancers. Complementarily, the temporal TF *hbn* is independently placed under spatial control in Mi21, highlighting the flexibility of patterning control of tsTF expression. It is important to emphasize that within the Vsx domain of the OPC, the same neuroblasts generate neurons that express *Vsx1/2* either through spatial mechanisms such as (early-born) Pm3 and Pm4, or through independent (temporal) mechanisms (typically later-born) such as Dm2^12,15^. The tsTF *bsh* was also found to be regulated by different upstream mechanisms in the medulla^37^ compared to the lamina^38,39^, and likely through different enhancers^40^; however, it is worth mentioning that lamina neurogenesis does not involve neuroblasts, nor do the progenitors undergo temporal patterning. The only other demonstrated case in *Drosophila*, to our knowledge, is the regulation of *knot* in VNC^41,42^, however the two neurons implicated in this case are generated from different NBs.

The orthogonality of the different progenitor patterning programs has long been appreciated in the *Drosophila* nervous system: in the developing VNC^21^, central brain^43,44^, and optic lobe (OPC)^14^, there are neuroblasts that undergo the same temporal cascade (in each region) despite being in different locations. There has been active debate about how these progenitors programs may instruct specific neuronal identity programs, especially because there appears to be little direct continuity in the expression of spatial and temporal TFs themselves in neuronal progeny^12^, and even less so in their enhancer-usage^7^. Recent work in the VNC^27^ and central brain^26,45^ proposed to extend this orthogonal framework of the progenitor patterning programs to the neuronal identity programs (terminal selectors), such that most of the specific tsTFs expressed by neurons are instructed either by their spatial/Notch (hemilineage) or temporal origin. There are several tsTFs in the OPC that are expressed in almost all neurons from one or a few consecutive temporal windows, ignoring most spatial boundaries. These are known as “concentric genes”^37^ and include Dll (**Fig. 2a**), Bsh, Drgx, Runt, Vvl, Lim3 and Ets65A^12^. However, it is more common to find tsTFs like Hbn, which is expressed in a domain-agnostic manner in a Dichaete temporal window that generates T1 neurons^12,13^; in the later Tll temporal window that generates Mi15/21 neurons, it is under restricted spatial control (**Fig. 6**). Unlike temporal patterning, in the medial OPC, there are no tsTFs that are expressed in all neurons sharing a particular spatial origin (and no others); only Vsx1/2 and Bifid have strong correlations that are far from perfect^13,15^. Overall, most terminal selectors appear to be regulated jointly by spatial and temporal patterning. For *Vsx1/2* and *hbn*, our results indicate a complex mode of regulation involving multiple enhancers responsive to different combinations of patterning factors. Their expression can be specified both by spatial and temporal patterning, even among neurons generated from the same NBs. We therefore favor a modular model in which spatial, temporal and Notch inputs combinatorially converge on different enhancers of largely overlapping sets of selectors, rather than being parceled out among dedicated tsTFs.

In multiple studies^12,18,37^, including this one, the OPC temporal TFs have been genetically linked to the expression of specific tsTFs. It is, however, not straightforward to unravel how the specification program (developmental origin defined by patterning regulators) instructs the initiation and maintenance program (terminal selectors) in each neuron. This is in part due to the transient nature of this process; spatial patterning factors are not expressed within NBs or GMCs^14,16^ and temporal TFs are minimally expressed at the RNA level in young neurons. This is exemplified by the regulation of *Vsx1* by BarH1: While we cannot yet be certain that this regulation is direct, we note that, in our OPC trajectories, the expression of these two genes never overlap at the RNA level (**Fig. 1c’-c”**). However, it is possible that like Tll in glia^18^ and M24 neurons (**Fig. 2c**), BarH1 protein inherited from progenitors is transiently present in newborn Dm2 neurons to initiate *Vsx1/2* expression. Our multiome data^7^ enabled us to identify temporal window-specific enhancers for *Vsx1/2*. By training ChromBPNet models that predict cell-type-specific accessibility directly from DNA, we were able to use their base-resolution contribution scores to nominate the functional TF binding sites in one of these enhancers. Deletion of these specific sites from a reporter construct sharply reduced its activity in Dm2 neurons. That a model given only accessibility, and no prior knowledge of Vsx biology, converged precisely on the sites that matter *in vivo* suggests that the instructions routing developmental origin to terminal identity are surprisingly legible in the DNA.

The redeployment of a spatial patterning factor as a neuronal identity regulator is neither unique to the optic lobe nor to invertebrates. In fact, the mouse homolog VSX2 (Chx10) was shown to function both in retinal progenitors and as an identity factor for bipolar interneurons^46^. These two roles are encoded by physically separable modules of a single super-enhancer^47^, and this modular architecture is conserved from mouse to human^48^. It is unknown whether the progenitor VSX2 is involved in specifying neuronal VSX2 expression, but our findings here would caution against such an assumption as a default regulatory mechanism. The context-specific deployment of a Vsx gene through separate cis-regulatory modules in these distant species indicates a common regulatory logic that has been conserved across more than 500 million years of evolution.

The regulation of *Vsx1* also raises striking parallels with the Hox family of genes, the canonical master regulators of rostrocaudal patterning in all bilaterians. They spatially pattern the neural tube during embryonic development and can transmit this identity by dedicated Polycomb chromatin domains to enforce their spatially-restricted expression in the corresponding neuronal progeny^49^. Recent findings indicate that Vsx1 also interacts with Polycomb Repressive Complex to restrict (most of) its neuronal expression to the corresponding spatial domain^31^. Thus, the spatial “memory” can survive from NE to neurons within the chromatin architecture, despite entirely distinct enhancers driving the same gene, and despite the fact that Vsx1 is downregulated in NBs before being re-activated in neurons^29^. In further similarity, the importance of postmitotic Hox expression has also been widely established, for example, for motor-neuron columnar and pool identity in the vertebrate spinal cord^50,51^, and for segment-specific neuronal survival and terminal identity in the *Drosophila* VNC^52,53^. Their continued presence is required to maintain identity even in fully mature neurons, indicating that they function as terminal selectors^54^. Finally, just like Vsx1, there is evidence that the neuronal Hox code need not faithfully copy the progenitor domain, even though it is often assumed to be a direct readout of it. Indeed, low-level Hox expression has been reported in restricted thalamic and sensory-cortical neurons of the adult mouse, territories classically considered Hox-free^55^. Together, these parallels suggest that re-deployment of a spatial patterning factor in postmitotic neurons through different enhancers that need not respect its progenitor domain is a recurrent evolutionary strategy.

What emerges is a model in which terminal identity is not passively inherited from the progenitor but computed in each newborn neuron through an active re-encoding of patterning inputs. A spatial patterning factor can be placed under temporal control, and a temporal factor under spatial control, through dedicated enhancers of the same terminal selectors, even among neurons born of a single lineage. Such inputs are therefore not partitioned across separate TFs but integrated combinatorially at specific cis-regulatory elements, each reading whichever signals its lineage and environment makes available. The observation that the same logic recurs from the *Drosophila* optic lobe to the mammalian forebrain^56,57^ suggests this is a general solution to the problem of converting a finite patterning code into the vast diversity of neuronal types.

## Acknowledgements

We would like to thank all members of the Özel Lab for helpful discussions and Tom Kleist for technical assistance. We thank Claude Desplan and Ted Erclik for both providing key reagents and critical reading of the manuscript. We also thank Josie Clowney, Robb Krumlauf and Filipe Pinto Teixeira Sousa for critical reading of the manuscript. This work was supported by the Stowers Institute for Medical Research and NIH R00-NS125117.

## Author Contributions

M.N.Ö. conceived the project. R.C., C.L., R.R. and B.T. performed experiments. R.C., F.K., R.R., M.T., Y.-C.C. and J.Z. analyzed data. R.C. and M.N.Ö. wrote the manuscript with input from all authors.

## Declaration of Interests

Authors declare no conflicts of interest.

## Data Availability

A published single-cell multiome dataset^7^ (GSE305940) has been used in this study. All original data underlying this manuscript can be accessed from the Stowers Original Data Repository [LIBPB-2646] upon publication.

---

**Supplementary Figure 1:**
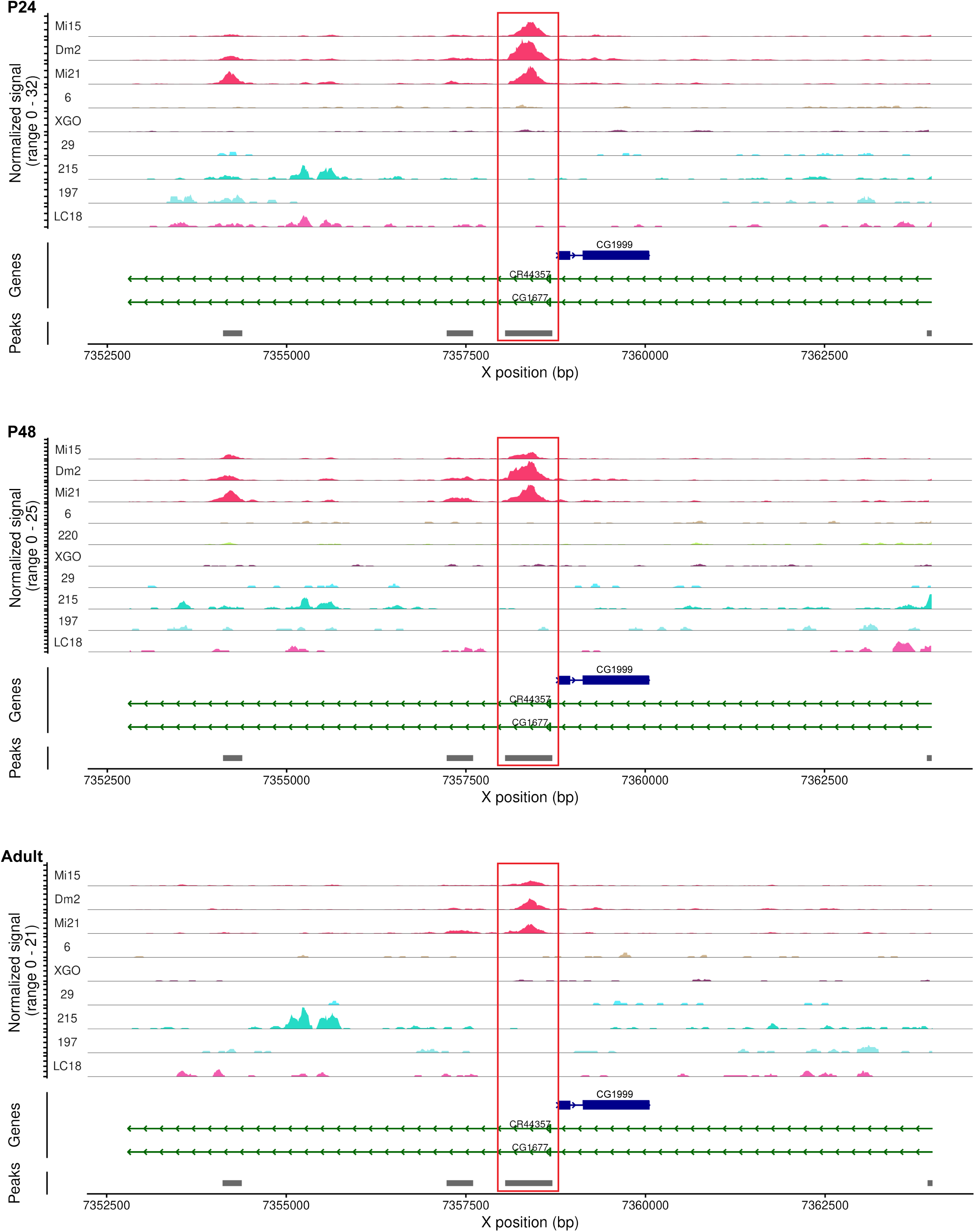
Accessibility of the enhancer region cloned as *M24-Gal4*. Coverage plots of the P24, P48, and Adult ATAC signal in each cell type showing normalized accessibility around the region that was cloned as *M24-Gal4* (red rectangle). The top 10 cell types (numbers represent unannotated clusters7) with highest accessibility at P48 are shown. Cluster 220 is only present at P48.

**Supplementary Figure 2:**
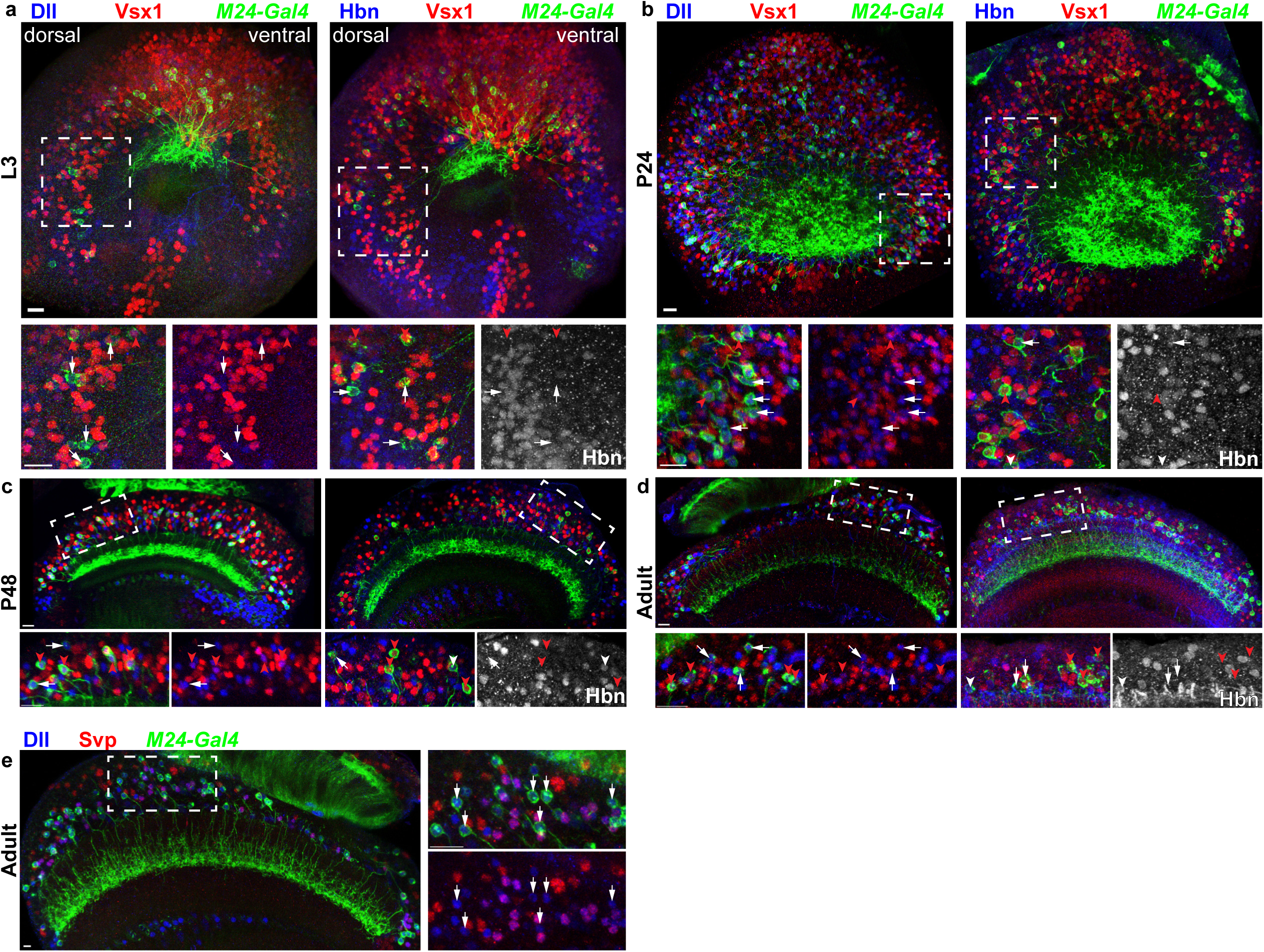
*M24-Gal4* expression during optic lobe development. *M24-Gal4* driving *CD4:tdGFP* expression across development (L3 in **a**, P24 in **b**, P48 in **c**, Adult in **d**-**e**). In **a**-**d**, left panels show anti-GFP (green), anti-Dll (blue), and anti-Vsx1 (red), with panels of higher magnification for regions marked by dashed white boxes below showing only Dll+ expression (white arrows) or Dll+Vsx1+ co-expression (red arrowheads) among the GFP-labeled cells. Right panels for each stage show anti-GFP (green) anti-Hbn (blue) and anti-Vsx1 (red) with panels of higher magnification for regions marked by dashed white boxes below showing a mixture of Vsx1+Hbn- (red arrowheads), Vsx1/Hbn+ (white arrowheads), and Vsx1-Hbn- (white arrows) cells. The delayed activity of *M24-Gal4* in Mi21 neurons (no expression at L3), along with the previously reported lower effectiveness of *UAS-Vsx2* compared to *UAS-Vsx1*25, likely explains the failure of *UAS-Vsx2* to convert Mi21 into Dm2 (Fig. 1d). In **e**, the left panel shows anti-GFP (green), anti-Dll (blue), and anti-Svp (red) with panels of higher magnification marked by dashed white boxes shown to the right. White arrows mark cells of Dll+Svp-. Magnified regions of the L3 brains are in the dorsal Optix domain. Representative images are shown as maximum intensity projections. L3, n = 3/5, P24, n = 3/4, and P48, n = 2/5 anti-Dll brains/anti-Hbn brains and Adult, n = 5/4/5 anti-Dll brains/anti-Hbn/anti-Svp brains. Scale bars: 10 μm.

**Supplementary Figure 3:**
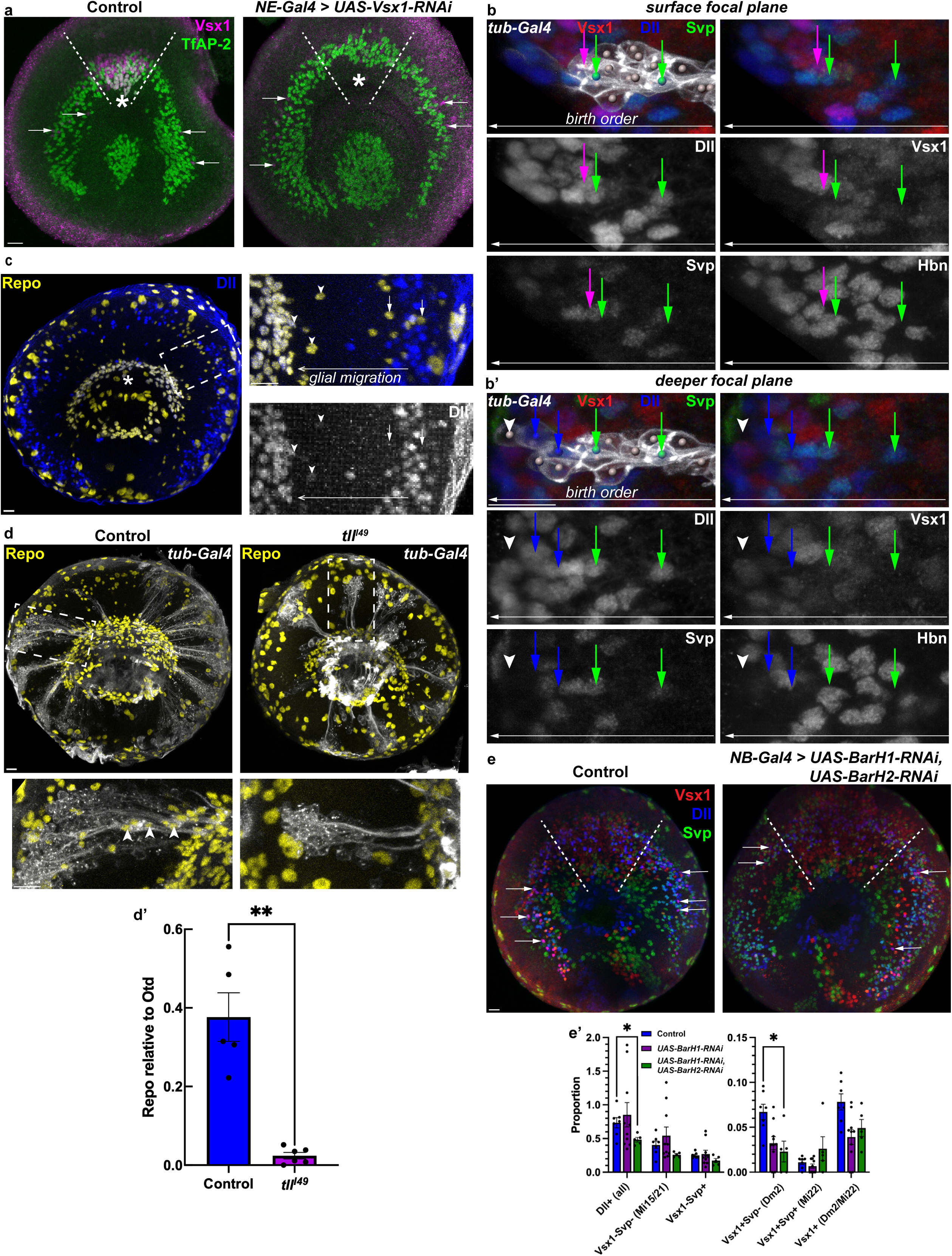
Birth order of Distal-less^+^ cells in the OPC and Tailless mutants. **a**, The same genotype as in Fig. 2a. Neuroepithelial driver (*R29C07-Gal4*) driving *UAS-Vsx1-RNAi* during neurogenesis (L3) with control lacking RNAi. Dashed white lines mark the Vsx1 spatial domain, asterisks mark TfAP-2+/Vsx1+ neurons produced exclusively from the Vsx1 domain and white arrows point to Vsx1+ cells outside the domain. Anti-Vsx1 (magenta) and anti-TfAP-2 (green) are shown as maximum intensity projections within a representative sample. n = 4/5 control/experimental biological replicates. **b**, The same clone shown in Fig. 2b. For clearer visualization, it was split into two focal planes (**b’**) with Dll+/Svp+ neurons common to both panels and Dll+/Vsx1+/Svp- (Dm2) and Dll+/Vsx1-/Svp- (Mi15) neurons in separate panels. FRT82B MARCM clones labeled with *tub-Gal4* driving *myr:GFP* in early pupal (∼P5-P10) optic lobes, approximately 50 hours after heat shock. Clones are located within the dOptix domain. Neurons are born from sequential temporal windows from right to left marked by the white arrow. All cells marked with arrows are Dll+ with Svp+ cells marked by green arrow and Vsx1+ cells by a magenta arrow. Anti-GFP (white), anti-Dll (blue), anti-Vsx1 (red), and anti-Svp (green) with single panels for each TF including anti-Hbn (white) that is not shown in the composite panel. n = 4 brains. **c**, A wild-type optic lobe during neurogenesis (P5) that shows the glia which originate from OPC NBs turning on Repo after their migration (arrows) to the neuropil edge (asterisk). Dll expression is stochastic and low within these glia while they migrate and increases as they reach the neuropil (arrowheads). Anti-Repo (yellow) and anti-Dll (blue or white) are shown as maximum intensity projections. The region within the white box is shown with higher magnification (right). n = 6 brains. **d**, The same genotype as in Fig. 2d, FRT82B (control) and FRT82B, *tllI49* MARCM clones are labeled with *tub-Gal4* driving *CD4:tdGFP* during neurogenesis (P5). 1-2 clones with arrowheads marking migrating medulla neuropil glia that express Repo are shown per genotype. Anti-Repo (yellow) and anti-GFP (white) are shown as maximum intensity projections with regions marked by white boxes shown with increased magnification (below). **d’**, Quantification of d. Ratios were quantified relative to the number of Otd neurons (*Methods*). Error bars represent SEM and each dot is one optic lobe with all clones summed. n = 5/5/26 control and n = 6/6/24 optic lobes/ brains/ clones. In comparison to the control, p = 0.0043 for Repo+ glia. **e**, Neuroblast driver (*R13C02(dpn)/NB-Gal4*) driving *BarH1* and *BarH2* RNAi during neurogenesis (L3) with control lacking RNAi (same as in Fig. 2e). Dashed white lines demarcate the Vsx spatial domain. Arrows point to Dm2 neurons (Dll+/Vsx1+/Svp-). Anti-Dll (blue), anti-Vsx1 (red), and anti-Svp (green) are shown as maximum intensity projections. Scale bars: 10 μm. **e’**, ROIs within both dorsal and ventral Optix regions were quantified and normalized to Otd (*Methods*). Error bars represent SEM and each dot represents one optic lobe. Only Dll+ cells were quantified. Quantification of the control compared to *UAS-BarH1-RNAi* are in Fig. 2e’. Adjusted p-values for each cell type are Dll+ (all), p = 0.0488, Dll+/Vsx1-/Svp- (Mi15/21), p = n.s., Dll+/Vsx1+/Svp- (Dm2), p = 0.0488, Dll+/Vsx1+/Svp+ (Mi22), p = n.s., Dll+/Vsx1+ (Dm2/Mi22), p = n.s., Dll+/Vsx1-/Svp+, p = n.s. for *UAS-BarH1-RNAi, UAS-BarH2-RNAi* in comparison to the control. n = 13/7/6 control, n = 15/10/9 *UAS-BarH1-RNAi*, and n = 7/5/5 *UAS-BarH1-RNAi, UAS-BarH2-RNAi* Optix regions/lobes/biological replicates.

**Supplementary Figure 4:**
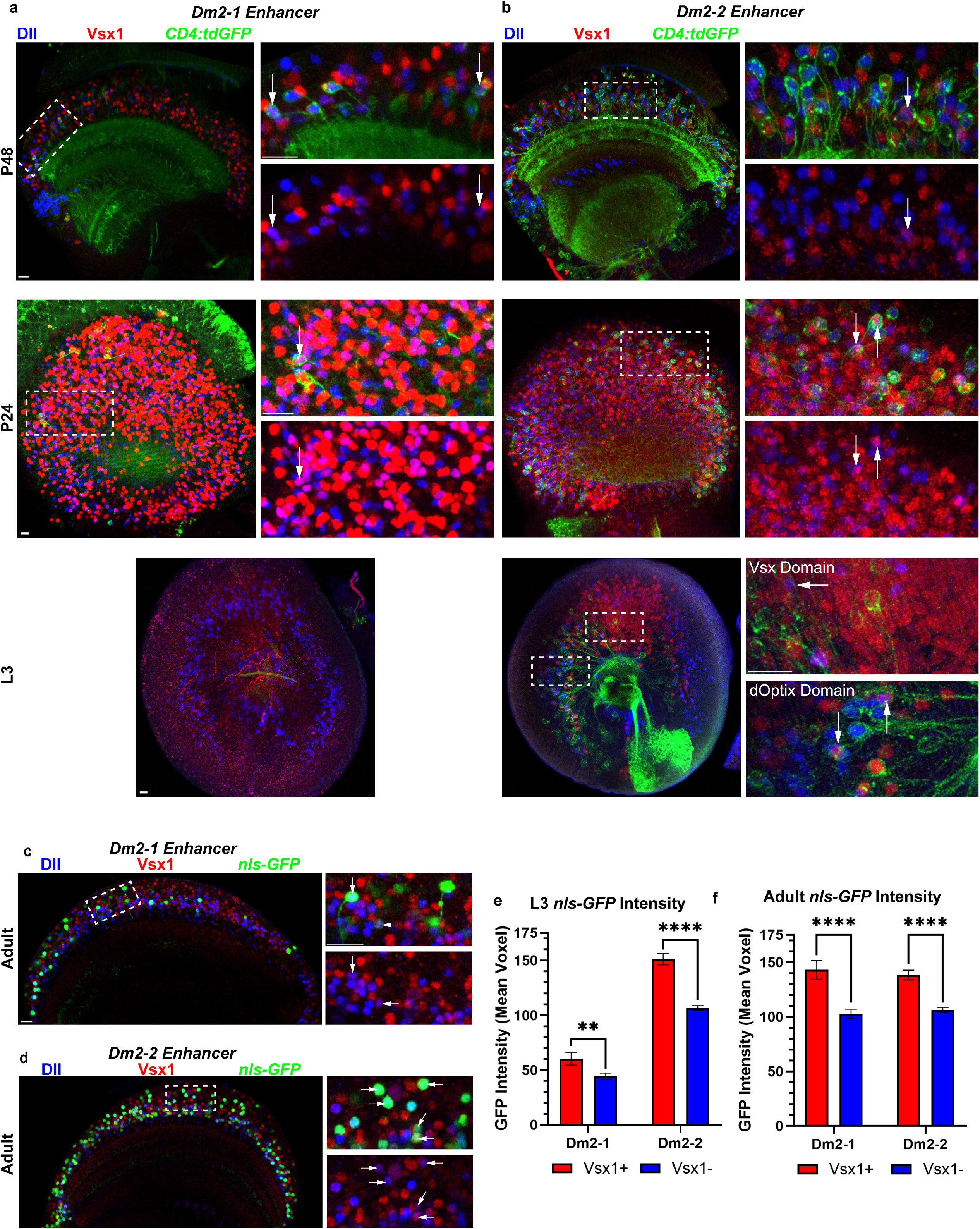
Dm2-1 and Dm2-2 enhancer reporter activity during optic lobe development. Dm2-1 (**a**) and Dm2-2 (**b**) enhancers were cloned into a *Gal4* reporter to drive *CD4:tdGFP* expression at stages P48, P24 and L3 and *nls-GFP* at the adult stage (**c-d**). Figure 4 shows the Adult stage with the same *CD4:tdGFP* and L3 with *nls-GFP*. Anti-GFP (green), anti-Dll (blue), and anti-Vsx1 (red) are shown as maximum intensity projections. White arrows point to Dll+/Vsx1+ co-expression within GFP labeled cells. Dashed white boxes indicate regions shown with higher magnification to the right of each image. For membrane labeling, P48, n = 4/2, P24, n = 4/4, and L3, n = 3/5 Dm2-1 brains/Dm2-2 brains. For *nls-GFP* at Adult (**c-d**), n = 5/5 Dm2-1 brains/Dm2-2 brains. Scale bars: 10 μm. **e**-**f**, Average nls-GFP intensity was measured in both Vsx1+ (red) and Vsx1- (blue) cells at both the L3 (**e**) and Adult (**f**) stages. For nls-GFP intensity measurements at L3 (**e**), n = 5/3 Dm2-1 brains/Dm2-2 brains. For comparisons between Vsx1+ and Vsx1- GFP intensities at L3, p = 0.00949/<0.0001 for Dm2-1/Dm2-2 GFP+ nuclei in brains. For nls-GFP at Adult (**f**, images in **c-d**), n = 5/5 Dm2-1 brains/Dm2-2 brains. For comparisons between Vsx1+ and Vsx1- GFP intensities at Adult, p <0.0001 Dm2-1/Dm2-2 brains.

**Supplementary Figure 5:**
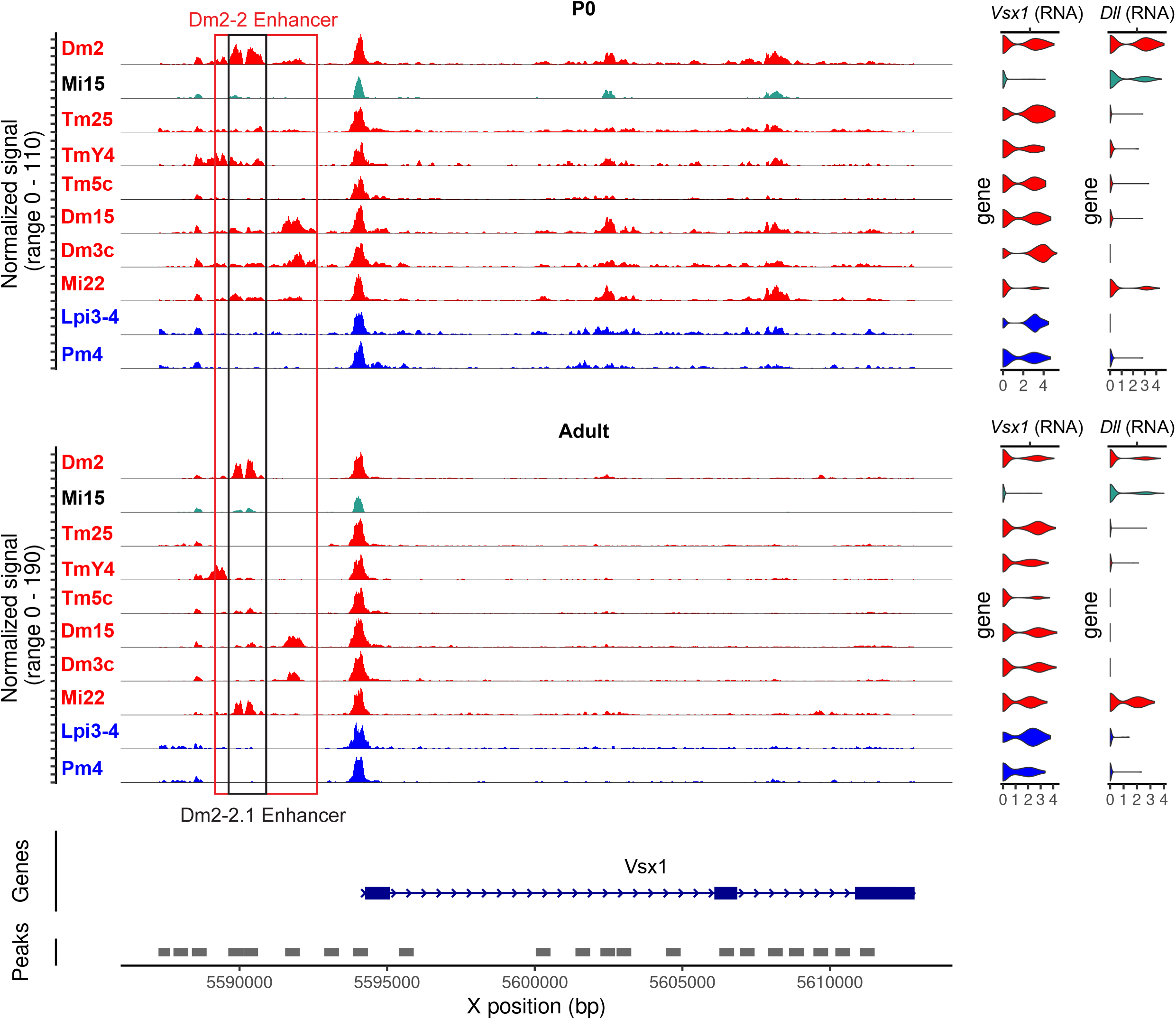
Dm2-2 enhancer accessibility in select cell types across development. Coverage plots of the P0 and Adult multiome data showing normalized accessibility around the ∼4kb Dm2-2 enhancer region in Dm2 and Mi15 neurons and additional *Vsx1/2*+ cell-types. At P0, the enhancer is broadly open in Dm2, while later the accessibility becomes restricted to distinct ∼1 kb subregions specific to each lineage (e.g., upstream in TmY4, downstream in Dm15/Dm3c, overlapping with Dm2 in Mi22). Violin plots (right) display log-normalized mRNA expression of *Vsx1* and *Dll* in these clusters. Lpi3-4 and Pm4 (blue) are representative cell types that originate exclusively from the Vsx domain and are spatially patterned while cell types in red originate from both Vsx and Optix domains.

**Supplementary Figure 6:**
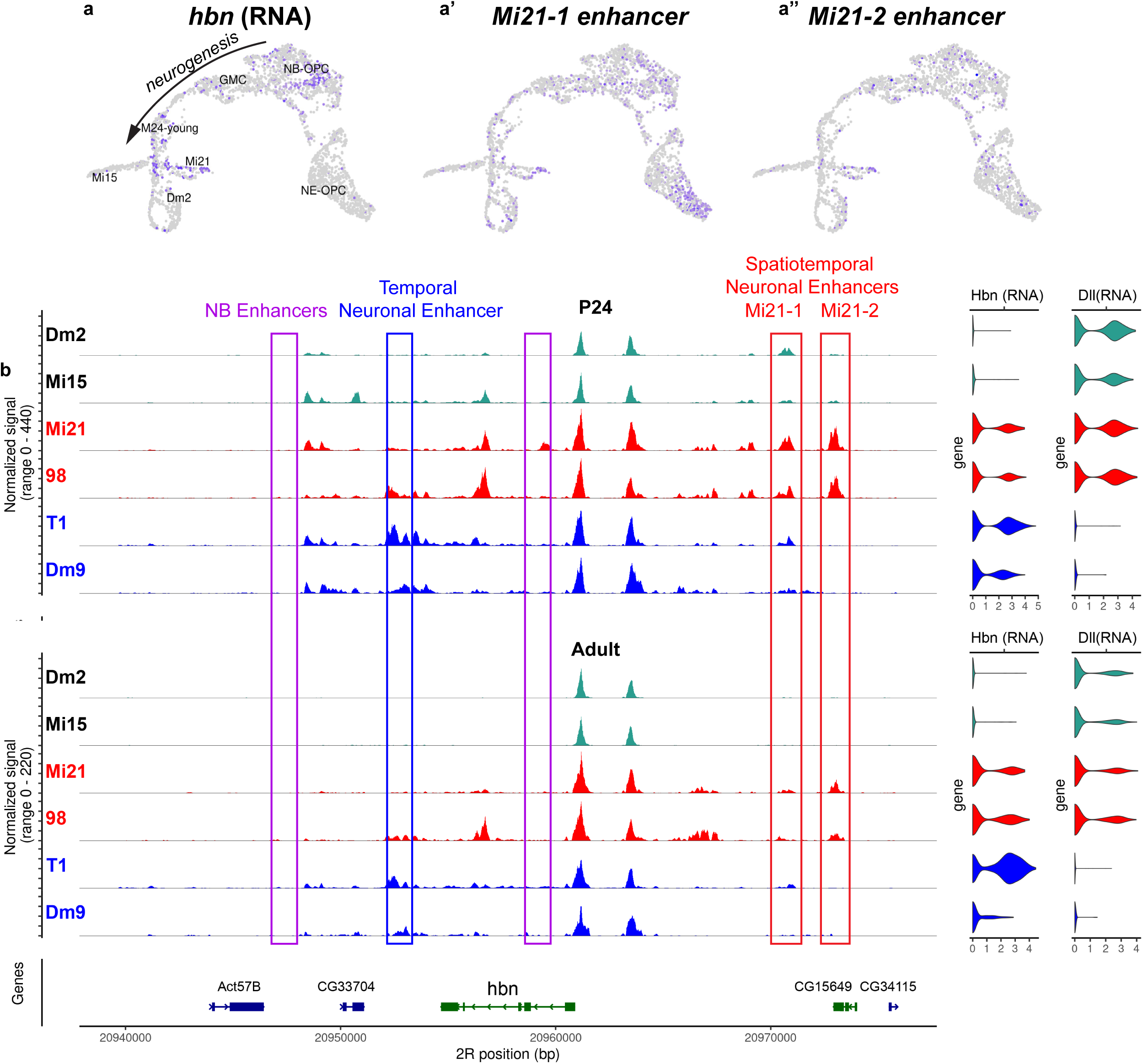
Accessibility of *hbn* locus across development. **a**, *hbn* expression, Mi21-1 (**a’**), and Mi21-2 (**a”**) enhancer accessibility displayed on the same UMAP as in Fig. 1c with cell types annotated. Color intensity corresponds to log-normalized and scaled RNA level (*hbn*) or ATAC signal (enhancers). **b**, Coverage plots of the *hbn* locus with tracks representing normalized accessibility in each cell type at P24 and Adult. Mi21 and cluster 98 (red) are late born cell types from ventral Optix domains. T1 and Dm9 (blue) are born from all spatial domains. Rectangles highlight possible cis-regulatory regions of NBs (purple), temporally regulated neurons (blue), and spatially regulated neurons (red). Log-normalized *hbn* and *Dll* mRNA expression in each cell type is shown on the right side.

**Supplementary Table 1:** Predicted transcription factor binding sites in the Dm2-2 enhancer. FIMO (find individual motif occurrences) analysis of the Dm2-2 enhancer (1.3 kb subregion, Dm2-2.1) identifying putative transcription factor binding sites. Only the binding sites for TFs differentially expressed among M24 neurons (Dm2, Mi15, Mi21) are shown. Each row reports the motif, predicted TF, strand, start and end coordinates within the enhancer, score, p-value, q-value, and matching sequence. These results highlight potential regulators, including a BarH1-like HD site.

## Supplementary Note

We reported that both Vsx enhancers (Dm2-1 and Dm2-2) we identified are active in Dm2 neurons as predicted, but the two reporters both labeled many other neurons, including some that were Vsx1^-^ (**Fig. 4c**). However, among the cells with detectable GFP expression, the intensity of the GFP signal (within a region of interest) was significantly higher in neurons that were Vsx1^+^ at both L3 (**Suppl. Fig. 4e**) and Adult (**Suppl. Fig. 4f**) stages. To test if the labeling of Vsx^-^ neurons is an amplification artifact of the Gal4/UAS expression system, we also generated a “direct” reporter for the Dm2-2 enhancer (**Fig. 4d**), where it directly drives the expression of membrane-tagged mCD4-tdGFP^58^. The relative proportion of Vsx^+^ cells labeled was higher in this reporter (**Fig. 4c,d**); it also labeled a significantly higher proportion of all Dm2s compared to *Dm2-2-Gal4* (73% vs. 48% in adults, p<0.0001). These results suggest that some, but not all, off-target (Vsx^-^) activity of these enhancers can be attributed to the reporter expression system and that cell types like Mi15/21 have some degree of activity (readout as reporter activity) that is not sufficient for driving *Vsx1/2* expression in the endogenous locus. This is reflected, to some extent, in the accessibility data as well: ATAC signal for the Dm2-2 enhancer in both Mi21 (**Fig. 3a**) and at later stages also Mi15 (**Fig. 3b**) is dramatically lower than what is observed in Dm2, but still visibly higher than the background levels seen in other neurons (e.g. the Vsx-domain neurons).

We can assume that this low (but clearly detectable) activity displayed by these enhancers in the reporter constructs is not functionally relevant in their endogenous locus, i.e. not sufficient to activate *Vsx1/2* expression. We suspect that this is a common feature of developmental enhancers, as it is consistent with the highly combinatorial nature of TF action. This finding implies that evolution must precisely tune the intrinsic strength of each enhancer, as its ultimate impact on gene expression will be a function of multiple other variables, such as the nature of each specific core promoter and the enhancer’s distance to it^59,60^, as well as the presence of shadow enhancers^61^ or silencer regions^62^ in the locus.

## Methods

### Trajectory analyses at P0

For the M24 trajectory in **Fig. 1c**, we took our multiome data^7^ and separated the NE-OPC and NB-OPC clusters, together with the P0 cluster 25 representing the late temporal window GMCs and the P0 cluster 13 containing newborn M24 neurons. Using MACS2^63^, we called peaks on fragments corresponding specifically to these barcodes. We then merged these peaks with the peaks we had determined for the M24 neurons from later stages. We recounted the ATAC modality of these cells based on these peaks and generated a new Seurat/Signac object. After standard processing steps and determination of PCA and LSI reductions, we used 20 PCA and 24 LSI components to generate the UMAP and cluster these cells. The annotations “M24-young”, “Mi15”, “Mi21” and “Dm2” in **Fig. 1c** were made based on these clusters, guided by the transferred labels from P24. The enhancers whose accessibility correlated with expression of *Vsx1* and/or *Vsx2* were determined using the LinkPeaks function; and their accessibility were further inspected on the UMAP and across different cell-types as described in Results.

### Generation of coverage plots

The genomic accessibility tracks in **Figs. 3,5** and **Suppl. Figs. 1,5,6** were plotted with *Signac*’s *CoveragePlot()* function using our published multiome data^7^ with the cell type annotations present in the *Seurat* object provided in GEO (GSE305940). The known subtypes of Pm3, Dm2, and Mi21 neurons were merged before plotting.

For the coverage plot in **Fig. 3a**, the neuroepithelial cells were separated into “Vsx” and “Other” domains by subclustering the original NE cluster. After subsetting the object, we calculated the top 30 PCA (RNA modality, SCT assay) and LSI (ATAC modality) reductions of the data using just these NE cells. We then inspected the correlations between the “loadings” of each of these reduced dimensions with the RNA expression of *Vsx1,* selected the top 9 PCA and LSI dimensions (each) with the highest absolute correlations, and calculated weighted nearest neighbors using the *FindMultiModalNeighbors()* function using only those dimensions. Clustering was then performed using this neighbor graph (resolution=2) to subdivide the NE cells. The subclusters were then assigned as “Vsx” or “Other” based on *Vsx1* expression.

Coverage plots of the Vsx1/2 locus (**Fig. 3**) were generated using Signac’s CoveragePlot() function on a merged object containing the NE, NB and GMC progenitor populations together with Dm2, Mi15, Mi21, and the Vsx-domain-exclusive neurons Pm3, MeSps, Lpi3-4, Pm4 and Dm21, with the ATAC assay set as default. The region X:5,560,183-5,594,255 was extended by 20 kb upstream and 10 kb downstream to include flanking sequence, and *Vsx1* mRNA expression was displayed alongside accessibility using the RNA assay (expression.assay = “RNA”, features = “Vsx1”). Tracks were colored by cell type using a custom palette.

For **Fig. 3b** and **Suppl. Fig. 5**, cell types Dm2, Mi15, Vsx-expressing cell types not from the domain (Tm25, TmY4, Tm5c, Dm15, Dm3c, Mi22), and Vsx-domain exclusive neurons (Lpi3-4, Pm4) were plotted from a merged object. The region of gene *Vsx1* was extended by 7000 bp upstream and *Vsx1* and *Dll* mRNA expression was displayed (expression.assay = “RNA”, features = c(“Vsx1”, “Dll”)).

For the coverage plot in **Fig. 6f**, the NBs were subclustered based on their age (Early, Mid, Late) as previously described^7^ and all GMCs were shown. For both **Fig. 6f** and **Suppl. Fig. 6**, cell types Mi15, Dm2, Mi21, cluster 98, T1, and Dm9 were plotted from a merged object. The region of gene *hbn* was extended by 15 kb upstream and downstream to highlight flanking regions and the mRNA expression of *hbn* and *Dll* was displayed (expression.assay = “RNA”, features = c(“hbn”, “Dll”))

### Animal Husbandry

All *Drosophila melanogaster* flies were reared at 25°C with males and females used for every experiment (except for FRT19A MARCM experiments which exclusively used females). Wild-type flies were either Canton S or *w^1118^*. Genotypes for every figure panel can be found at the end of these Methods with the origin of each stock reported in the Key Resources Table. Pupal dissections were staged by collection of P0 white pupa and waiting for 5 (for P5), 24 (for P24), or 48 hours (for P48) with vials kept at 25°C. L3 stage dissections were performed on large, wandering L3 larvae. Adult dissections were performed within 24 hours of pupal eclosion. Neuroblast MARCM clones were heat shocked at ∼50 hours for 10 minutes at 37°C before dissection at early pupal stages (P5-P10). *tll* MARCM flies were heat shocked for one hour at 37°C four days before wandering L3 were formed, consistent with previous experiments^12^. For both *spalt* and *disco* MARCM experiments, flies were heat shocked for one hour at 37°C three days before L3 formed. For all other MARCM experiments, wandering late L3 were transferred to a new vial and heat shocked at 37°C for 30 or 45 minutes. For *dpn-Gal4* driving *UAS-BarH1-RNAi*, vials were reared at 25°C for 3-4 days and moved to 29°C three days before dissection at late L3 stage. For experiments with a *Gal4* driving *UAS-CD4:tdGFP* (FLP-out), P0 were heat shocked for 5 minutes at 38°C. Sample sizes for each experiment are listed within figure legends. Animals were always selected randomly after confirmation of genotype.

### Immunohistochemistry

Animals were anesthetized on ice and then were dissected in cold Schneider’s Insect Medium with fixation in 4% PFA at room temperature for 25-30 minutes for pupal and adult brains with L3 brains fixed for 20 minutes. All wash volumes are 500 µL. Brains were briefly washed with 0.3% PBS-Triton (PBST) three times before nutating for 15 minutes. After washing, samples were incubated with primary antibodies at 4°C overnight. After the removal of the primary antibody solution, brains were washed three times for 10 minutes in PBST before incubation with secondary antibodies at 4°C overnight. The samples were then washed for 10 and 15 minutes in PBST and finally, for 15 minutes in PBS before mounting in Slowfade Gold with the desired orientation. All samples were imaged with a Leica Stellaris 8 confocal microscope using a 63x glycerol objective (NA=1.3). All antibodies used, their concentrations, and their origin can be found within the Key Resources Table.

### Image Analysis and Presentation

Images were created and analyzed within Imaris 9.6.1 and adjusted for figures within Adobe Illustrator 2025 and Adobe Photoshop 2025. They are shown as either maximum intensity projections to show protein expression within cells or with three-dimensional (3D) reconstruction using Blend mode with glow color settings to visualize more detailed neuronal morphology as indicated within figure captions. All quantifications were performed manually with assistance from Spot creation for cell body expression to aid in accurate and consistent counting. In sparsely labeled experiments like 19A MARCM, all cells were counted. Dm2 membranous reporters were quantified by creating ROIs that spanned the entirety of the Z stack where the signal was high enough for reasonable expression determination. At L3, all reporters were quantified with one ROI per domain (dorsal Optix, Vsx, ventral Optix). At Adult, Dm2-1 had one ROI centered on the more labeled side of the brain; *Dm2-2-Gal4* and its direct GFP reporter as well as all Dm2-2.1-Gal4 constructs had two ROIs to account for any variation that were then averaged for each sample.

For *Vsx1* knockdown within the neuroepithelium, *UAS-Vsx1-RNAi* was driven by neuroepithelial *R29C07-Gal4*. To quantify Dm2 within the Optix domains, ROIs of comparable sizes were centered on late Dll clusters within the Optix domains. Spots were created for Dll, Vsx1, and Svp with quality manually adjusted to represent real signal. Consistent spot settings for every channel were: Enable Region Of Interest = true, Process Entire Image = false, Enable Region Growing = false, Enable Tracking = false, Enable Classify = false, Enable Region Growing = false, Enable Shortest Distance = false, Estimated XY Diameter = 2.29 µm, Estimated Z Diameter = 3.50 µm, Background Subtraction = true. To measure co-expression, the spot of the weaker antibody was used to determine an appropriate threshold of mean intensity for the stronger antibody. Specifically, Dll spots were used to determine the mean intensity of Svp that was real signal and therefore determined Dll^+^/Svp^+^ co-expression. Similarly, Vsx1 spots were used for Dll and/or Svp co-expression. Dm2 (Dll^+^/Vsx1^+^/Svp^-^) was divided relative to total Dll as a representative ratio. *tll* perturbation in *tubulin-Gal4* clones were individually counted if Otd expression was present in that clone. Otd is a tsTF that is only expressed in neurons from the earlier Hbn temporal windows^12^, which should not be affected by loss of *tll*. Every combination of Dll, Vsx1, and Svp was quantified and summed across clones to create a single data point per optic lobe. For Repo quantification, only Otd and Repo cells were quantified. Because of the variation in sparsely labeled data, every sample in every experiment was quantified relative to other cells present to account for these variations and create a single data point per optic lobe.

For *BarH1* perturbation, *R13C02-Gal4* was used to drive *UAS-BarH1-RNAi* or *UAS-BarH1-RNAi; UAS-BarH2-RNAi* within all neuroblasts. To quantify the effect of the RNAi, regions of interest (ROIs) of appropriate size for each fully visible Optix domain of newly born neurons were created. ROIs include the entirety of late Dll while excluding earlier Svp expression and central brain Otd expression. ROIs were limited to maintain these standards for each brain. Within each ROI, spots were created for four channels (Otd, Dll, Vsx1, and Svp) with quality manually adjusted for each to represent real signal. Spots settings and co-expressions were created as described previously. All metrics were divided by Otd as a normalization with an identical assumption for the *tll* perturbation where Otd is a tsTF only expressed in neurons from the Hbn temporal window and therefore are not affected. Metrics were determined on a per lobe and per Optix domain basis. For each optic lobe, ventral and dorsal quantifications were summed and a single value per optic lobe was considered for a t-test because while the differing Optix domains had slight differences, they consistently had the same metrics as significantly different.

Mean GFP Intensity was measured for both Dm2-1 and Dm2-2 enhancer *Gal4* reporters as they drove *nls-GFP* within both Adult and L3 brains. For each brain, a sample ROI was created that excluded that most saturated neurons near the surface of the brain. Spots were created for the GFP channels with the following consistent parameters: Enable Region of interest = true, Process Entire Image = false, Enable Region Growing = False, Enable Tracking = false, Enable Classify = false, Enable Shortest Distance = false, Source Channel Index = 1 (GFP), Estimated XY Diameter = 2.25 µm, Estimated Z Diameter = 3.50 µm, Background Subtraction = true. For each sample, a rational Quality filter was applied to include as many real spots as possible while excluding non-real labeling. Within the spots, the mean intensity of both Dll and Vsx1 were observed to pick a rational threshold for ‘real’ expression which was later used to categorize the spots as either Dll^+^ and/or Vsx1^+^ for each sample. All mean intensity statistics for GFP, Dll, and Vsx1 were exported for comparison. Within L3 samples, the Optix and Vsx domains were distinct due to variability in staining.

### Statistics

For both *BarH* and *Tll* perturbations where all cell types were normalized to Otd expression (**Fig. 2d-e** and **Suppl. Fig. 3d-e**), parametric, two-sided Welch’s t-tests were performed at the optic lobe level if normal according to Shapiro-Wilk test (R package: rstatix, function: shapiro_test). If not normally distributed, Wilcoxon Rank Sum tests were performed. In analyses with multiple comparisons, Benjamini-Hochberg test was used to adjust p-values which are reported in figure captions along with sample sizes. For quantification of proportionality data like cell types labeled within *M24-Gal4* or *Dm2-enh1* and *Dm2-enh2-Gal4* reporters, quasibinomial (R package: stats, function: glm) generalized linear models (GLMs) were used. Quasibinomial GLMs are used to fit data with overdispersion, which we frequently observed due to biological variability between replicates. Within **Fig. 1g**, we observed complete separation where *hbn^15227^* mutants never produced Mi21 neurons which our quasibinomial GLM could not tolerate and produced infinity outcomes; we instead used a bias-reduced GLM (R package: brglm2) which yielded stable, non-infinity outcomes for Mi21:M24 ratio quantifications and applied this model to all cell types. For all analyses, individual optic lobes were considered as biological replicates within a genotype except for **Fig. 6c-d** where individual brains (i.e. two optic lobes) were taken as replicates due to very sparse labeling in those samples. All graphs were generated using Prism GraphPad with the lack of comparison indicating non-significant differences and the stars representing the following: * (P ≤ 0.05), ** (P ≤ 0.01), *** (P ≤ 0.001), and **** (P ≤ 0.0001).

### Creation of FRT19A, *Vsx1^A23^Vsx2^ΔEx3^*

*Vsx1^A23^* allele ^29^ was recombined with FRT19A. Within this stock, CRISPR-mediated mutagenesis by homology-dependent repair (HDR) was performed (by Well Genetics) to create a deletion of the third exon of Vsx2, which should remove most of its homeodomain in addition to creating a frame shift, resulting in a double mutant.

Two gRNAs, Cas9, and a dsDNA donor plasmid were injected into a strain of FRT19A, *Vsx1^A23^*/*FM7*, *Act>GFP*. sgRNA1 of sequence TTGAGATGGCTGCGAAGAGC[GGG] and sgRNA2 of sequence GCCAACAGCAAACACCTACG[CGG] were used, targeting a 688 bp fragment of Vsx2 that includes exon 3 (X:5,546,600-5,547,287), and integrating a cassette of GMR-RFP for screening. F1 progeny marked by RFP were validated by PCR and sequencing. The RFP cassette was flanked by LoxP sites, allowing for excision through Cre/LoxP recombination, which was validated again by PCR and sequencing.

### Enhancer Reporter Generation

The plasmid, pBPGUw, was a gift from Gerald Rubin^64^ (Addgene #17575). It was digested with EcoRI and FseI with the 1730bp containing attR sites, CmR, and ccdB between them removed and replaced with a multi-cloning site of sequence:

### GAATTCTTAATTAAGCTAGCAGATCTCCTAGGGGCCGGCC

We hereby refer to this new plasmid as pBPGUw-MCS. To create enhancer reporters, we amplified the corresponding regions with PCR performed on *Drosophila* genomic squashes (Canton S) and cloned them into pBPGUw-MCS. Successful clones were verified via Sanger sequencing using primers of AATAGGCGTATCACGAGGC (forward) and CTGATGCTCTCAGCCACCCC (reverse). The cloned plasmids were sent to BestGene for injection into BDSC Stock #8622 for PhiC31 integration into the attP2 site on the third chromosome. Lines were validated by the presence of the mini-white gene within the *w^1118^* background and were balanced over *TM3, Sb*.

Both Dm2 enhancers were identified by their significant peaks (**Fig. 3a**) and buffers of 100-200 bp were considered for primer selection. 25 mM DNA oligos were ordered from IDT. Dm2-1 enhancer was PCR amplified out of Canton S strain flies while Dm2-2 enhancer was PCR amplified out of DGRP-40 flies. Dm2-1 Enhancer was amplified from the (-) strand using primers of GCCGAATTCGTGTACAAGAATGATGGGCG (forward) and TATGCTAGCGTAGGCTGGACAAGTGAGCG (reverse) with final coordinates of X:5541310-5544572. Dm2-2 Enhancer was amplified from the (+) strand using primers of ATAGCTAGCGTGGCCCGACTCCATTTGCC (forward) and CCTAGATCTCAGCCCACAAGGCCAACCGATC (reverse) with final coordinates of X:5588668-5592926.

Restriction enzyme cloning (EcoRI-HF to NheI-HF for Dm2-1 and NheI-HF to BglII for Dm2-2) was used for insertion into pBPGUw-MCS. Both enhancers had multiple SNPs, INDELs, and/or polyN inconsistencies in comparison to the reference genome (*dm6*), but all PCR amplifications generated were identical, indicating no technical error.

The plasmid, pDEST-HemmarG2, was a gift from Chun Han^58^ (Addgene #112814). It was digested with SphI and KpnI with the 1736bp containing attR sites, CmR, and ccdB Gateway RfA cassette between them removed and replaced with a multi-cloning site of sequence:

### GCATGCCAATTGATCTGAAGAGCAGATCTGCGGCCGCGCTCTTCAGCCGGCCGGCCGGT ACC

We refer to this new plasmid as pDEST-Hemmar-G2-MCS. To generate a direct reporter for Dm2-2, the same bp with altered restriction enzymes were used to PCR the fragment from the previously made reporter plasmid. It was amplified using primers of ATAGCATGCGTGGCCCGACTCCATTTGCC (forward) and CCTAGATCTCAGCCCACAAGGCCAACCGATC (reverse). Restriction enzyme cloning using SphI and BglII was used to insert the same Dm2-2 fragment into pDEST-Hemmar-G2-MCS for a direct reporter with the same coordinates of X:5588668-5592926.

To generate the *M24-Gal4* driver, we analyzed the M24-specific ATAC peakset together with matched RNA data across developmental stages (P48, P24, and Adult) using Seurat v4.1.4 and Signac v1.13.0. Subtypes with nearly identical accessibility profiles (Dm2a/b and Mi21a/b) were merged prior to analysis (Dm2ab, Mi21ab). At each stage (P48 → P24 → Adult), cluster-specific peaks were identified sequentially using *FindMarkers* (test.use = “LR”, latent.vars = “atac_peak_region_fragments”, only.pos = TRUE, log2 fold-change > 2, min.pct > 0.05) in a one-versus-all manner for each M24 cluster (Mi15, Dm2ab, Mi21ab). Following each stage-specific search, peak lists from the three M24 clusters were merged and reduced to retain only peaks commonly enriched across all subtypes, yielding a progressively refined set of M24 candidate peaks at each stage.

The final Adult-refined list was then re-tested at P48 using FindAllMarkers with a lower stringency (log2 fold-change > 0.25, min.pct > 0.05, max.cells.per.ident = 500). This global comparison ensured that candidate regulatory elements were not spuriously detected as markers in non-M24 cell types and allowed us to quantify any off-target clusters that may show up.

The resulting peak sets therefore represent regulatory elements consistently and uniquely accessible in M24 cell types relative to all other neuronal and non-neuronal clusters across development. Marker peaks were further annotated with peak–gene linkage scores using Signac’s *LinkPeaks* function (±100 kb from transcription start sites) and assessed for overlap with validated enhancer collections from Janelia and Vienna Tiles. Candidates were then plotted and visualized using Signac’s *CoveragePlot* function, and this peak was chosen manually. (**Sup. Fig. 1**). This chosen candidate was cloned from the (+) strand using primers of TCTGAATTCTTTTCTTGGCTGTCAGCGAGC (forward) and TCAGAATTCCCCTTCGCCTTTAATGTCTAG (reverse) with coordinates of X:7357748-7359076. Note that both primers have an EcoRI-HF site allowing for integration in both directions within the plasmid and the forward (5’ to 3’) was chosen via sequencing. Following the same method as described above, the fragment was amplified from Canton S, cloned into pBPGUw-MCS, and injected by BestGene for PhiC31 integration at the attP2 site.

To create a Dm2 specific enhancer reporter of Dm2-2, which we call Dm2-2.1, the broader Dm2-2 region was restricted to X:5589618-5590901. The wild-type sequence cloned in *Dm2-2-Gal4* was copied exactly and synthesized at GenScript.

Wild-type sequence of Dm2-2 Dm2 specific region is GAATTCTTTTGTAGCTAAACAGAACGACACAAATTATTCTTAATCTCAAACAATTTGGAAATA AATTAATTAGATTTACAAGCATATTTTGCGTTAGAAAAGAATACATTTCTGCTTAACCCCCTGC TCCTTTTTCTTAACCCTTTTCCCACTTCTTTTTTTGGTACCAACCCTTCGAGTCAAAGAAATC TAAAAATGCAAGAGAAAGAACCCAAGTGGCCGCGTGGAAGGGGCTTATTTTTCGAAAGGA ATGGAGCAGAAGTTGACCAATTATGTAAACCACTTACACCACTGTTACATATGTATATGTACC TCACACACACAAACACACGTAAACGCACACGCACACAACCTCGCAGTGGGGGGAAAAAAA ACTGAGAACGAAAACACGTCAATGCGATAATTATTAATAATTATCCACTCGTCCTTCTTTTGC TTCTTCTGTTGCCGCTGCCCCTGTTTCTGCCTCTTTCTACCACTGCTTCCCTCTCTCTCCC ACTACTTCTCACTTGCAGCACTGAGAGAAAAACGTATCATAAAGATTTAAATACTGAACTGA ATCTAAATCCATCTAAATTTCAGTTCTAATTATCAGTAATAATGATAGAAATGCATAAGTTATTA TCTTAGAATAAAGTCTGAAGATTAAAAACACAACTAGATCATTTTCGCGATGTGTAGCTGGC TTGTTGTGACCGCTCCTTCGAGGCCACTTCGTCCACATCGAGGCGCTCCCTGGCCACCA GTCCACCTCCCTCCATGCGAATCCTGGTGCTGATTTCCCGCTGAGTGGCATTGCAACAAA GGCCCCCCCCCCCCGCTCCGCCAACAAGTTCGTGTGGCCTGAACTGCTAATGGACGGGC AAAACTAAATTATGGCCAAAATGGTGTTGGCTTTCTGGCTGGCTGCCACTTATGTGCCACT CACACATCCACAAAACGTCCGCAGCGACACATACACACATCCCATATGTAAATAAGCGTATA TATATATATCTGAATATACATATATTGTAGTACAACCCGGACTTCCCCATTTTGGAGTTCGGAG TTCGTAGCGTGTGCATGAAACATGTAATTATGAAATTAACGATTTCCAAGTGGCTCAAAGCG ACGTCGCCATTTGGGAGCGTGTCAGCCAGTCAACCAGTCAGCCAGGATAACGCAAAGGA ACGGCAGAACAATGGCCAGGATAATGTGGCGCAAGGATGCACAAGTGTCGGTTTGGGTAC AGTCTGTATTTTGAGAGTTCATCTCAAATGTTGCAAACCCCTTCCGCTGATTAGCTGCTGCT TTTGCTAGC. Flanking restriction enzymes sites of EcoRI and NheI were used to insert the fragment into pBPGUw-MCS.

FIMO was run with default parameters using differentially expressed TFs within the M24 metacluster with all HD motif sites tested within our ChromBPNet models. The PWM database was downloaded from CisBP (v2). Motifs were named 1-6 from 5’ to 3’ as they occurred and are annotated within **Suppl. Table 1**. There were many SNPs between the reference Dm6 sequence and what was cloned from DGRP-40; only one SNP was within any HD motif. Motif HD_5 had a T>A where TTTAA<u>T</u>TAC became TTTAA<u>A</u>TAC. Dm2-2.1ΔHD346-Gal4 had the following regions deleted: CGTAAACGCA, CAATGCGATAATTATTAATAATTATC, and GTAATTATGAAATTA. Dm2-2.1ΔHD1-6-Gal4 had the previous motif deletions and these additional: ATAAATTAATTAG, ACCAATTATG, and TTTAAATAC. As described above, plasmids were injected by BestGene for PhiC31 integration at the attP2 site.

### ChromBPNet

Model training was performed using a BPNet-style deep learning framework implemented with BPReveal^65^. For each selected cell type and developmental stage, ATAC-seq bigWig tracks from the optic lobe developmental multiome dataset^7^ were used as chromatin accessibility signal and paired with consensus peak coordinates from the same dataset. Although the original dataset included many cell types, the analyses here were restricted to the Dm2 and Mi15 cell types.

Candidate regions were partitioned into training, validation, and test sets (80/10/10), with final regions filtered by signal intensity. To account for sequence-specific Tn5 insertion bias, we first trained a bias model using an approach analogous to ChromBPNet^35^. This bias model captured Tn5-associated sequence preferences without learning transcription factor motifs, allowing the full accessibility models to learn residual accessibility not attributable to enzymatic bias. Cell type-specific models used a first convolutional layer with a 7-bp filter receptive field, eight dilated convolutional layers, 256 filters per layer, a 31-bp profile head, and a 1,000-bp output window. Models were optimized in a joint profile-and-counts framework using a batch size of 128, a learning rate of 0.002, early stopping, and learning-rate plateau monitoring. All training jobs were executed on GPU nodes through a SLURM-based workflow with a fixed random seed to improve reproducibility. Model performance was evaluated on held-out test regions, and fold-level Spearman correlation between the predicted and observed counts was used as the primary criterion for retaining models for downstream interpretation. Five models were generated for each of the Dm2 and Mi15 cell types from the P0 dataset, with counts spearman correlations of 0.68, 0.66, 0.73, 0.71, 0.72 for Dm2 models and 0.66, 0.68, 0.68, 0.65, 0.67 for Mi15 models.

To interpret the trained models, we generated predictions for peak-centered input regions and computed nucleotide-level contribution scores to identify sequence features predictive of chromatin accessibility. Model interpretation was performed using a deepSHAP^36^-style attribution framework, in which contribution scores are calculated with respect to the predicted accessibility counts for each input sequence. These nucleotide-resolution attribution values were summarized as contribution tracks and used to identify sequence positions with the strongest influence on model predictions. Contribution scores were derived for each model fold for all peak regions, and the mean computed across folds. The averaged predicted accessibility profiles and contribution scores were then converted to bigWig files and used for all analyses and figures.

### Genotypes of experimental strains

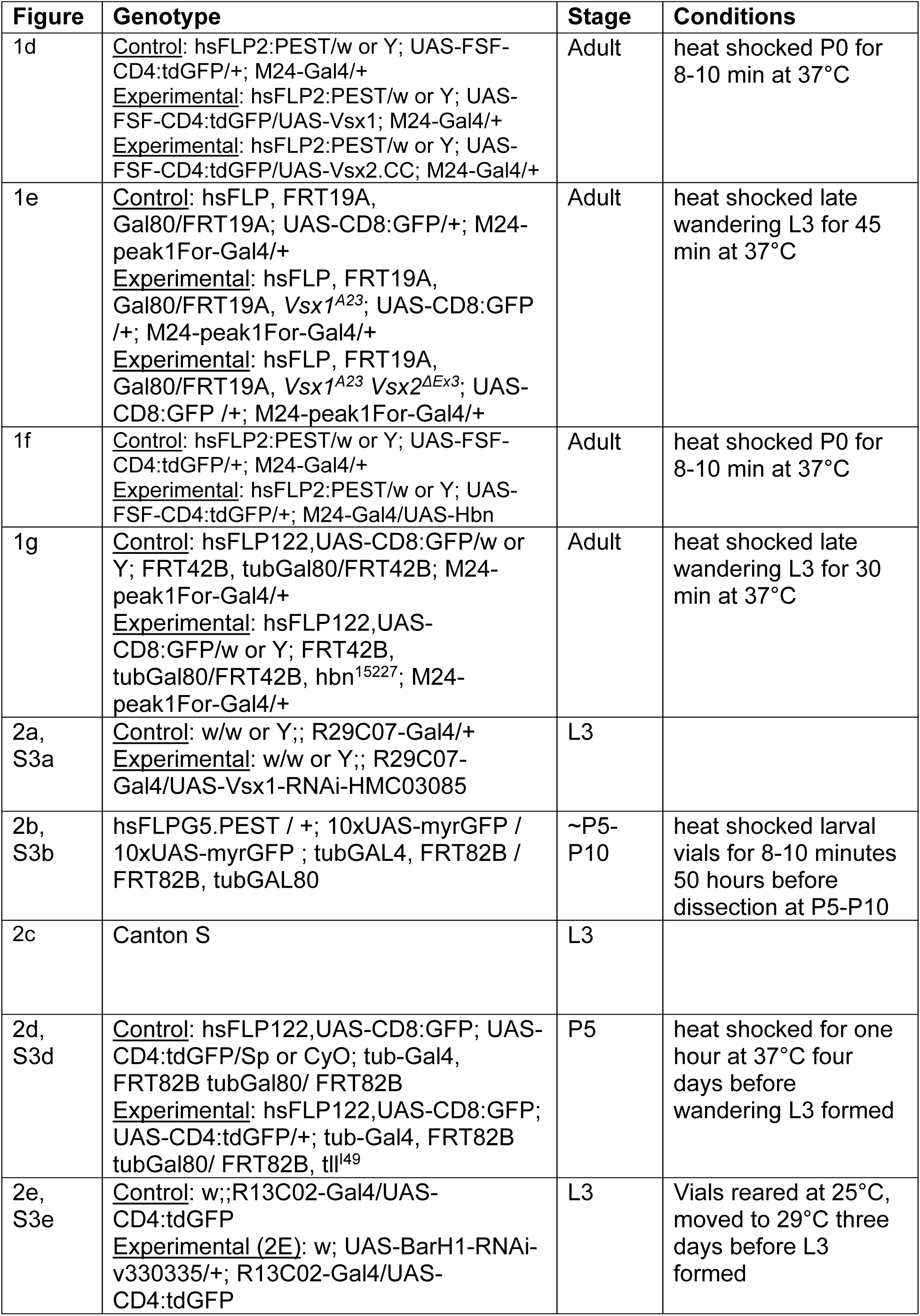
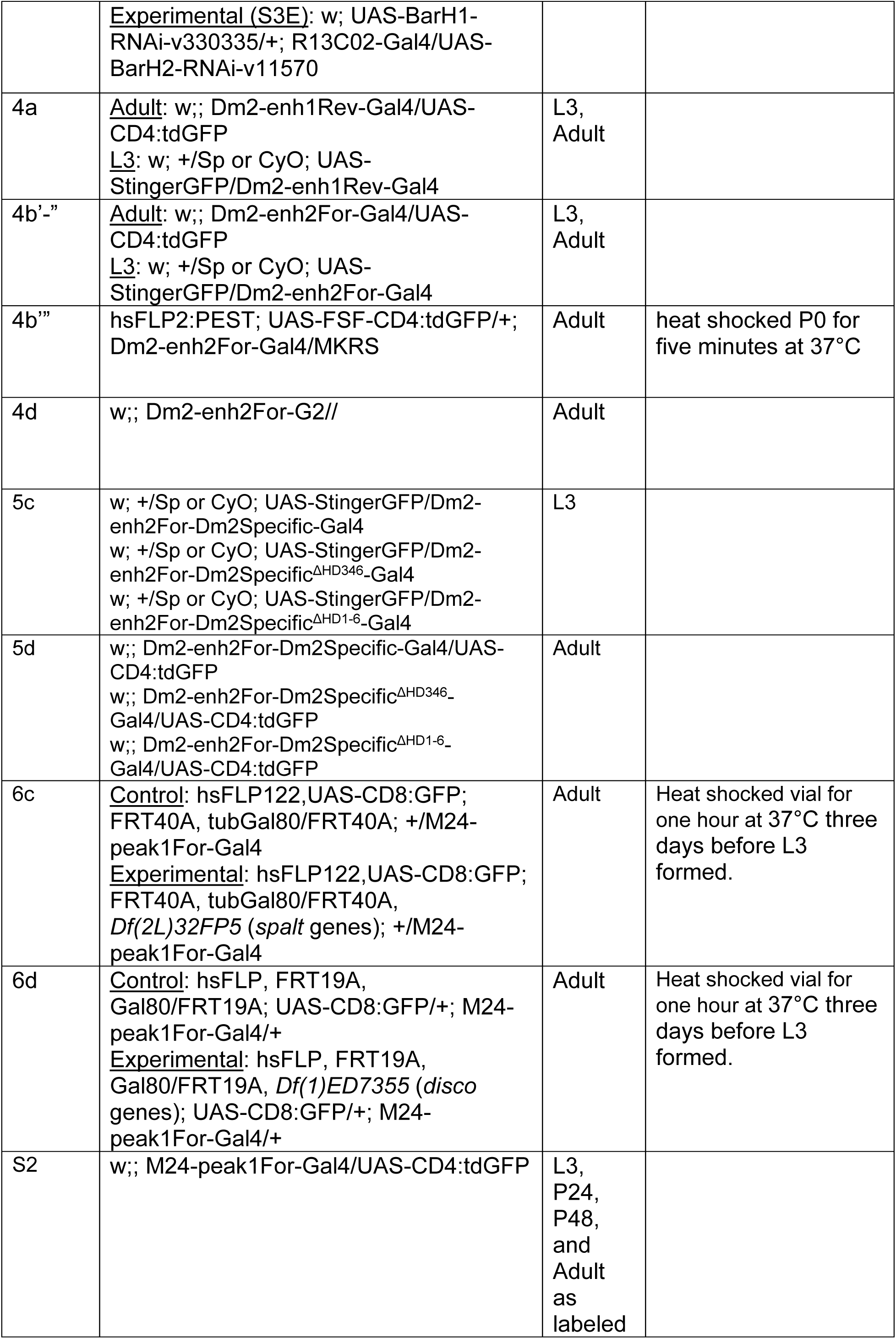
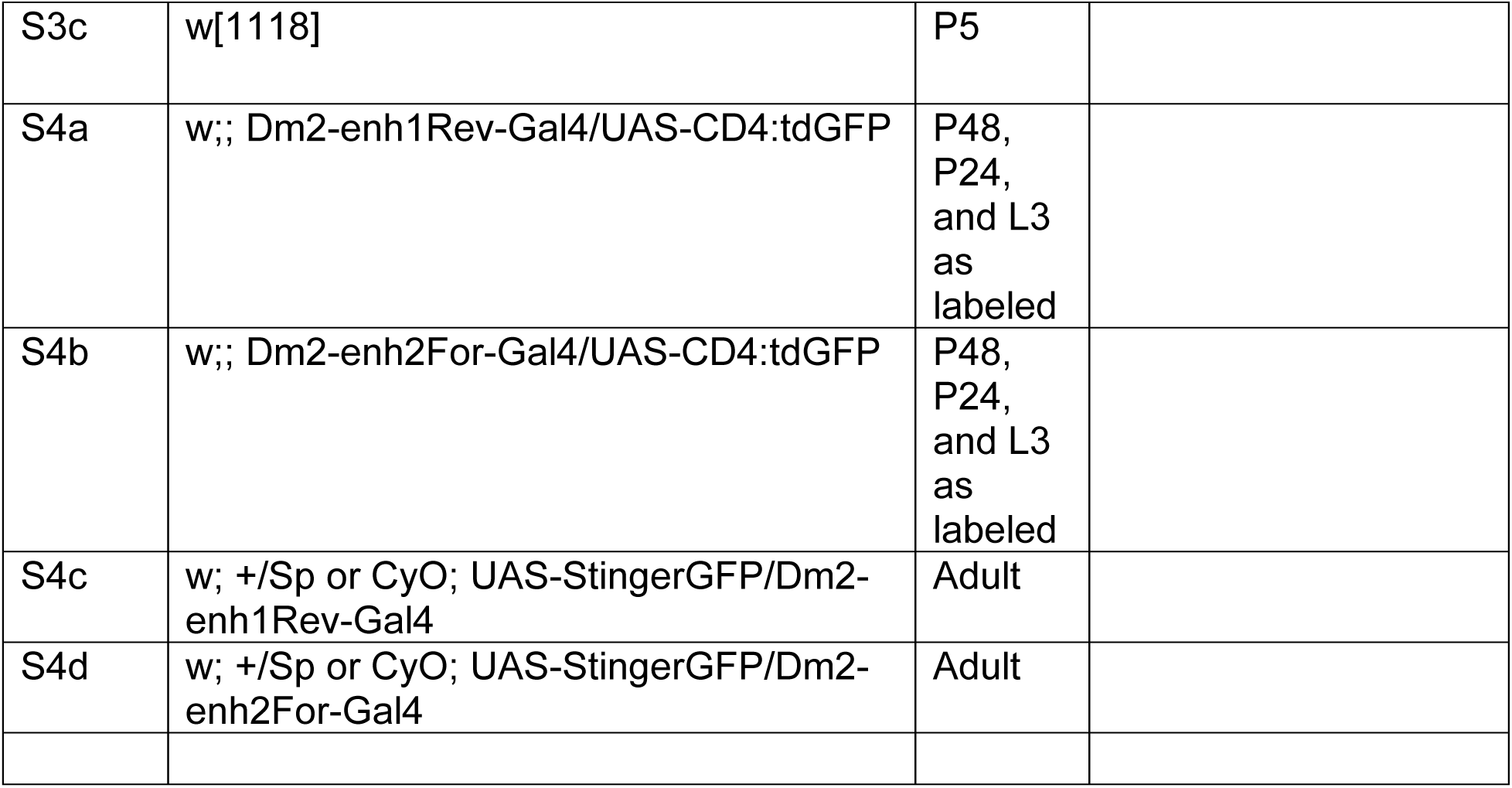

### Key Resources Table

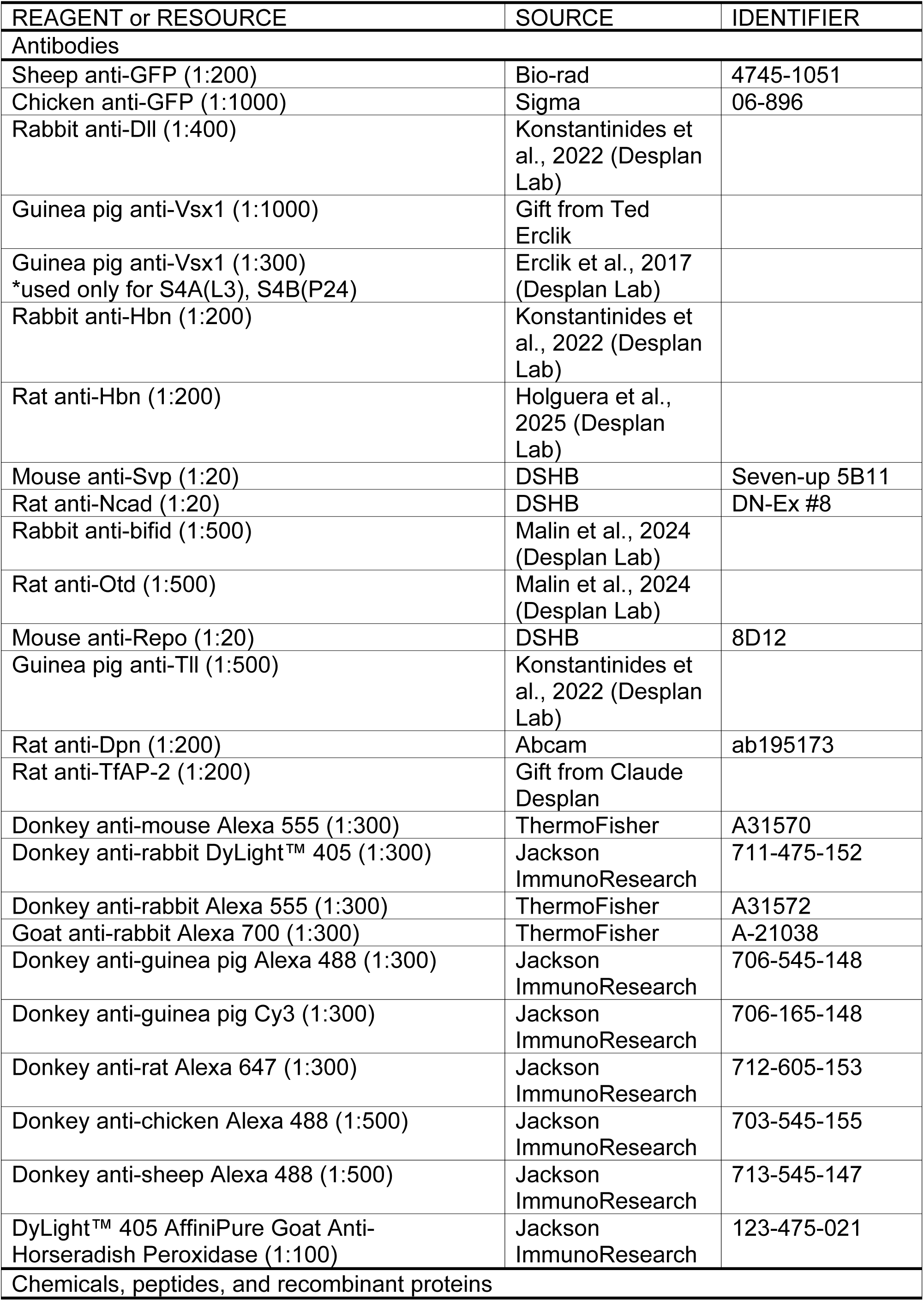
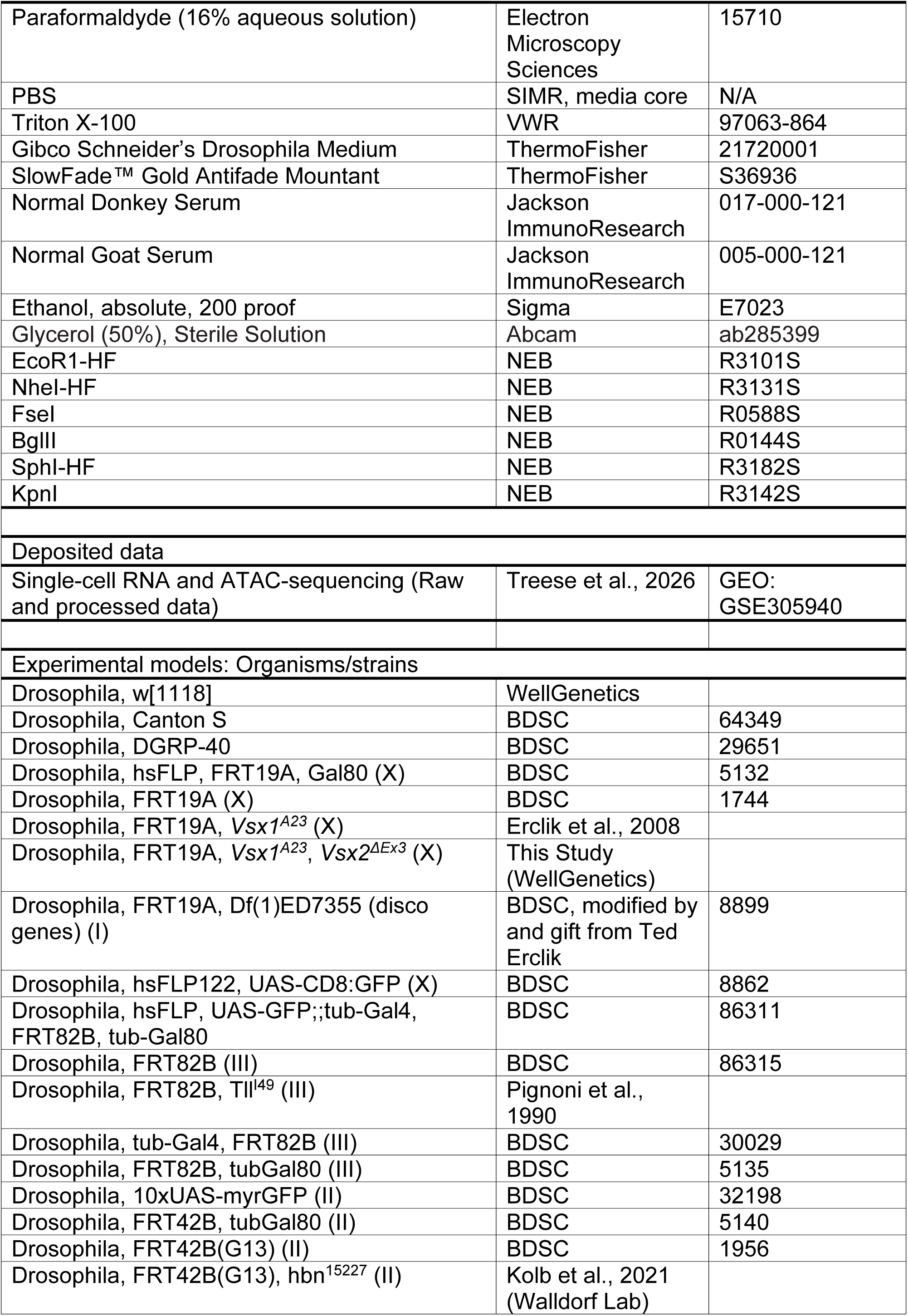
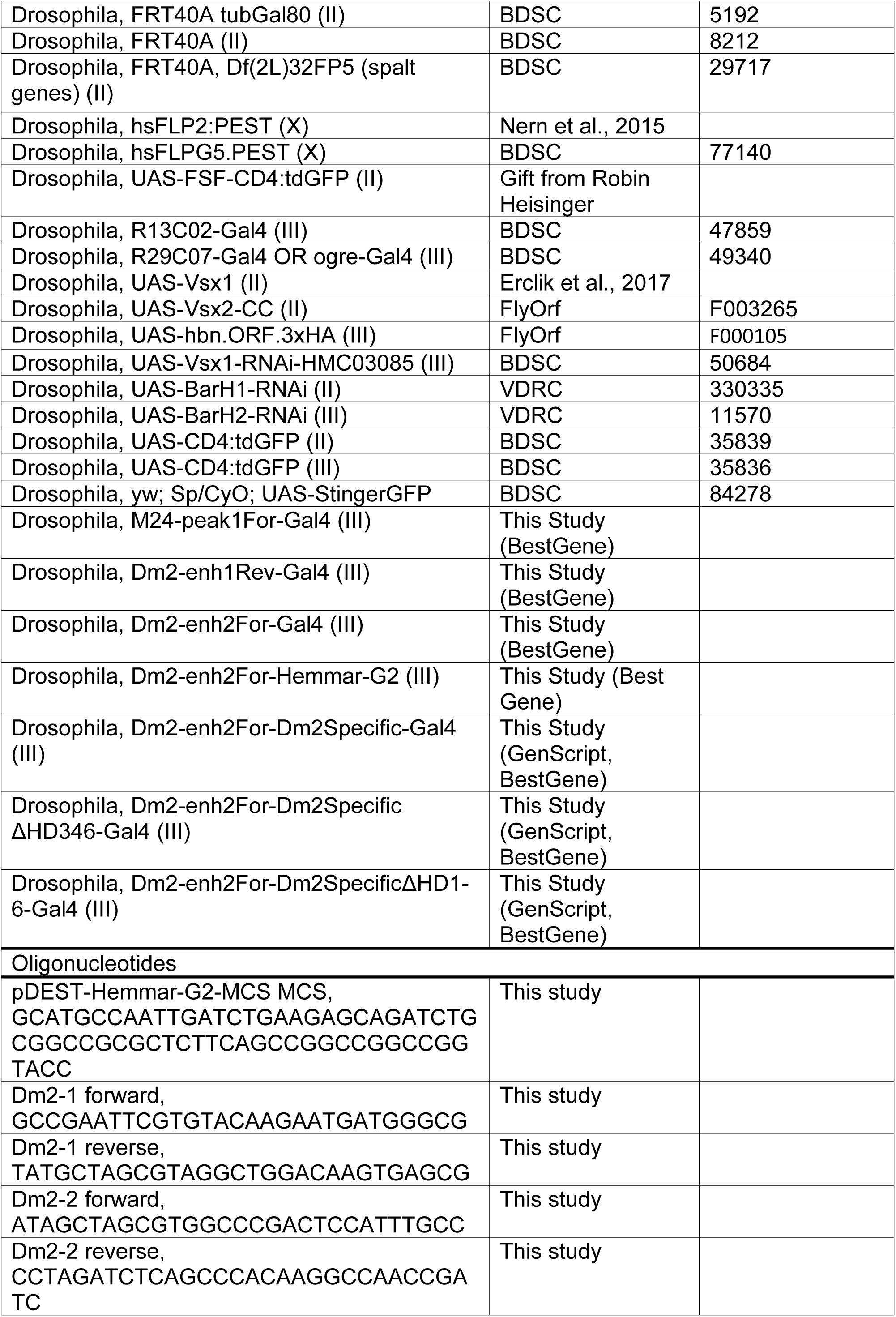
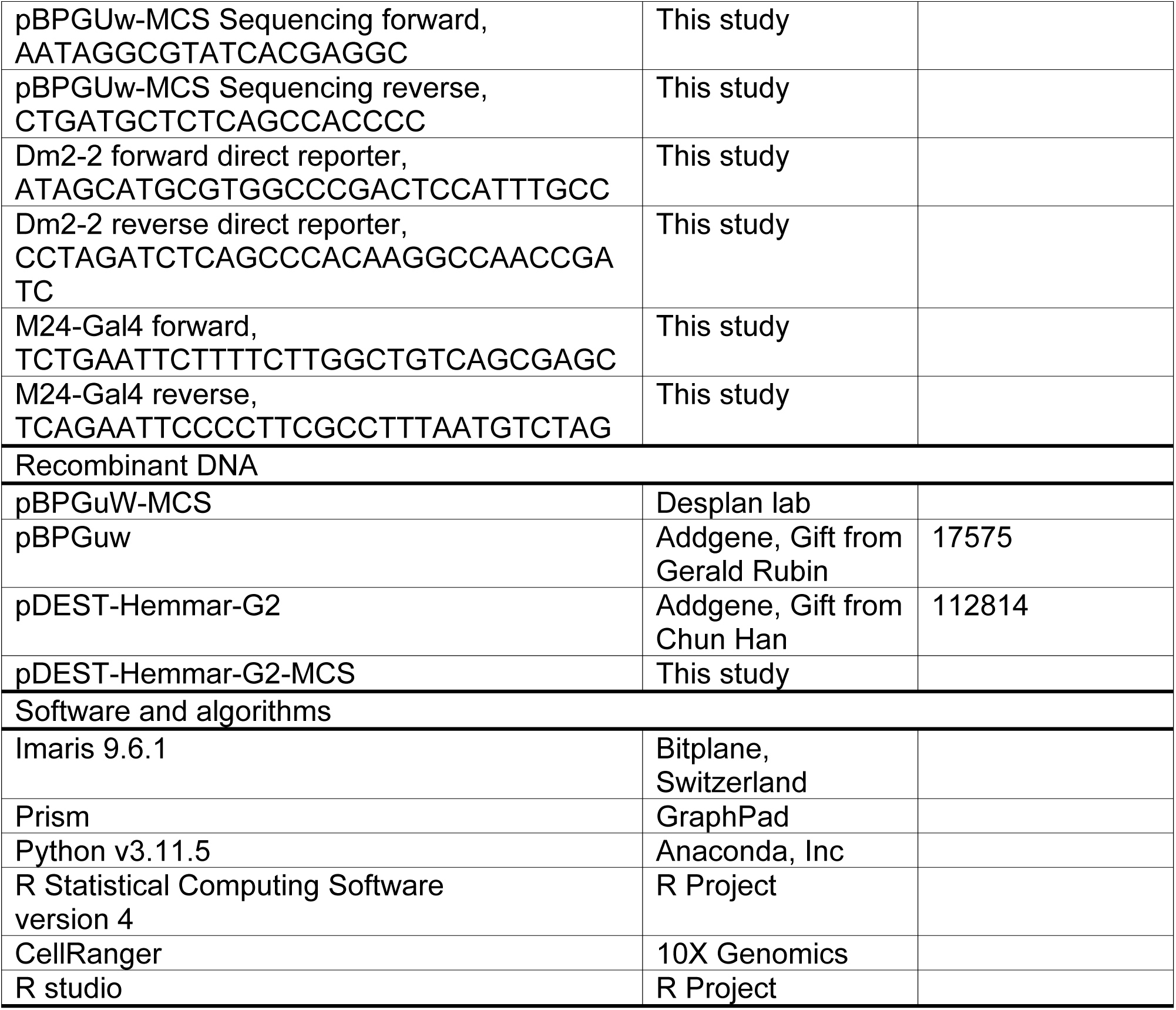

## Notes

### Competing Interest Statement

The authors have declared no competing interest.

### Summary of Updates

Minor corrections and reorganizations. /

